# Unlocking subcellular imaging of a cnidarian photosymbiont *Breviolum minutum*, through expansion microscopy

**DOI:** 10.64898/2026.08.19.745656

**Authors:** Pranali Deore, Cameron J. Nowell, Volker Leen, Douglas R. Brumley, Madeleine J. H. van Oppen, Elizabeth Hinde, Johan Hofkens, Linda Blackall

## Abstract

A cnidarian photosymbiont alga, *Breviolum minutum*, is an emerging model to study symbiosis due its ability to colonise host in absence of light, and amenability to genetic and physiological manipulations. This alga undergoes subcellular reorganisation in response to stress conditions such as elevated temperature and nutrient deprivation. However, subcellular visualisation of this alga is challenging because of its broad spectrum autofluorescence (400–700 nm) and relatively small size (6–8 µm). We developed a super resolution imaging, Expansion Microscopy (ExM) workflow – a hydrogel-based technique for mechanical enlargement of cells, that reveals previously inaccessible subcellular features in *B. minutum*. This ExM workflow presents a set of thermic and enzymatic conditions which enables 4-fold expansion of *B. minutum*, optical clearing of autofluorescence as well as the removal of its thick cellulose rich cell wall. We implemented a recently described platinum (II)-based tri-functional linker 1, to retain *in situ* hybridised oligonucleotides targeted to 18S rRNA within ExM hydrogel and exploited its azide reactive group for post-ExM fluorophore labelling (DBCO modification). We observed actin patches (a cytoskeletal feature) and calmodulin (a calcium binding signalling protein) that are not previously visualised in *B. minutum*. This approach overcomes some of the long-standing problems in visualisation of *B. minutum* using commonly available reagents and commercially available low-cost ExM compatible chemistries. The broader uptake of this tool for the visualisation of diverse species of photosymbionts will pave the way for fundamental discoveries underpinning cellular reorganisation in formation and breakdown of symbiosis.

## Introduction

A marine dinoflagellate, *Breviolum minutum* (formerly clade B in the Symbiodiniaceae family) forms close association with marine prokaryotes (Maire et al., 2021) and cnidarian hosts (Nitschke et al., 2022). This alga is emerging as a model to study symbiosis due to its ability to colonise host in photosynthesis dependent or independent manner (Jinkerson et al., 2022) as well as its amenability to genetic (Gornik et al., 2022) and physiological (Kirk et al., 2020) manipulations. Symbiotic associations governed by *B. minutum* and symbionts alike are at brink of collapse due to climate change related stressors (Hughes et al., 2018). Extensive efforts are underway to investigate how a range of climate driven stressors such as elevated temperature, altered nitrogen levels (Pasaribu et al., 2016), and low pH (Ishii et al., 2023) impact subcellular organisation in dinoflagellate symbionts. Prolonged suboptimal growth conditions, especially elevated temperature, in cultured photosymbionts lead to changes in their cell size (Amario et al., 2023; Rosic et al., 2024), membrane fluidity (Oakley et al., 2022; Tchernov et al., 2004), antioxidant pool (Majerová and Drury, 2022), relative distribution of photosystem I and II embedded within thylakoid membrane (Deore et al., 2024b; Slavov et al., 2016), biomacromolecule composition (Lima et al., 2022), membrane lipids (Tortorelli et al., 2025) and microbiome (Camp et al., 2020). However, visual confirmation of these subcellular changes in *B. minutum* using light microscopy has remained challenging due to the presence of broad spectrum autofluorescence spanning from 450–750 nm of the visible light spectrum (Deore et al., 2024b), a thick cellulose rich cell wall (Pairs et al., 2024), and a relatively small cell size (6–8 µm for *B. minutum*) (Nitschke et al., 2022). This limits the use of traditional immunofluorescence techniques combined with conventional confocal laser scanning microscopy (CLSM, 200–300 nm resolution), as well as advanced light microscopy modalities such as Stimulated Emission Depletion (STED, 20–30 nm resolution) for specific and/or non-specific labelling of sub-cellular targets in *B. minutum* (Deore et al., 2022). Therefore, innovative imaging modalities overcoming these challenges for subcellular imaging of *B. minutum* are urgently needed.

In our previous work, we explored fluorescence lifetime imaging microscopy (FLIM) modality for the distinction of autofluorescence, arising from photopigments, from intracellular bacteria labelled with a fluorescent probe in *B. minutum* (Deore et al., 2022). The FLIM approach relies on the measurement of fluorescence lifetime of fluorophores instead of their intensity which is commonly mapped using CLSM. The fluorescence lifetime gating approach helped eliminate autofluorescence signals in *B. minutum* (Deore et al., 2024a, 2022), however, FLIM setup requires a specialised hardware combined with CLSM which is not commonly available in most laboratory setups. The relatively small cell size (6–8 µm) of *B. minutum* (Nitschke et al., 2022) combined with the resolution limitation of CLSM microscopes further hinder the observation of closely spaced sub-cellular targets which, in part, have also impacted the use of spatial omics tools (van Oppen and Raina, 2022) for the study of *B. minutum*. A readily accessible super resolution imaging modality combined with optical clearing of autofluorescence is vital to advance our efforts in mapping of sub-cellular organisation in *B. minutum*.

In this work, we hypothesise that an implementation of a hydrogel-based technique, expansion microscopy (ExM) (Chen et al., 2015), will aid in optical clearing of autofluorescence arising from endogenous chemicals (Valdes et al., 2024; Wassie et al., 2019) and mechanical enlargement (Chen et al., 2015) of cultured *B. minutum* cells. Further, we expect the combination of ExM with specific (FISH, antibody) (Wen et al., 2021), non-specific (NHS ester) (M’Saad and Bewersdorf, 2020), along with click chemistry (Sun et al., 2021) labelling approaches utilising trifunctional linkers will enable visualisation of subcellular features that were previously not easily observed in *B. minutum*. We outline detailed optimisation of ExM procedure (thermal and enzymatic conditions) to achieve consistent 4-fold isotropic expansion of *B. minutum* cells using commonly available chemicals. Using this procedure, we distinctly observed pyrenoid, F-actin patches and a calcium binding protein, calmodulin in *B. minutum* without any autofluorescence. Overall, our robust ExM will pave the way for the detailed mapping of diverse subcellular features (cytoskeleton, mitochondria, photosystem I and II, etc.,) and nuclear architecture of *B. minutum* that will advance our knowledge about this dinoflagellate’s stress responses.

## Materials and Methods

### Breviolum minutum cultures

*Breviolum minutum* (SCF127-01, Australian Institute of Marine Science, Townsville, Australia) was cultured in Daigo’s IMK medium (1% w/v, NovaChem, Heidelberg, Australia) prepared in filtered Red Sea^TM^ salt water (fRSS, Reefs Secrets, Burleigh, Australia) of 34 practical salinity units (PSU). Cultures were incubated at 26°C under a 12 h:12 h light:dark cycle with 40–50 µmole per m^2^ per s light intensity (model no: 740FHC LED light chambers, Taiwan Hipoint Corporation, Kaohsiung, Taiwan). Exponential phase cultures (2 mL) were harvested at 1 x 10^5^ cells per mL and centrifuged at 10,000 *g* for 5 min. Supernatant was discarded and the cell pellet was treated with a different combination of chemical fixatives as outlined in sample fixation step of ExM.

### Reagent preparation and storage

1. Chemical fixatives: Paraformaldehyde (PFA, catalogue no. 15710, Emgird, Para Hills West, Australia) solution was diluted to 8% (v/v) in fRSS and stored at 4°C until use. Acrylamide (AA, catalogue no. A9099, Merck, Buchs, Switzerland) solution was prepared by dissolving 4 g of powder in 10 mL of milliQ water (40 % v/v) and stored in 4°C for up to six months. Ready to use formaldehyde solution (FA, catalogue no. F8775, Merck, Milwaukee, Wiconsin, USA) of 36.5–38% was commercially procured and stored at room temperature (RT).
2. FISH hybridisation buffer: The hybridisation buffer was freshly prepared by mixing 20 mM of Tris-HCl (pH - 7.4), 0.9 M of NaCl, 0.01% sodium dodecyl sulphate (SDS) and 25% formamide in milliQ water.
3. FISH wash buffer: The wash buffer was freshly prepared by mixing 20 mM of Tris-HCl (pH 7.4), 0.149 M of NaCl, 0.01% SDS and 40 mM of ethylenediaminetetraacetic acid (EDTA) in milliQ water.
4. Permeabilization buffer: Triton^TM^ X-100 (catalogue no. A16046.AE, Thermo Fisher Scientific, Haverhill, Massachusetts, USA) was dissolved (0.2%, v/v) in 3X Gibco^TM^ phosphate buffer solution (PBS, catalogue no. 10010023, Paisley, Scotland, UK) and stored in 4°C until its use.
5. Cell wall enzyme digestion mix: Cellulase RS (4%, w/v) and macerozyme R-10 (1%, w/v) powders (catalogue nos. C8003.0001 and M8002, respectively, Duchefa Biochemie, Netherlands) were slowly dissolved in fRSS and filtered using 0.45 µm Millex^TM^ syringe filters (catalogue no. SLHVR33RB, Merck, Darmstadt, Germany) to remove any undissolved matter. Aliquots (2 mL) were made in centrifuge tubes and stored in −20°C for only up to three months.
6. Poly-L-Lysine solution (catalogue no. P8920, Sigma-Aldrich, Missouri, USA) was diluted to 0.01% (v/v) in milliQ water and stored in 4°C until its use.
7. Coverslip and glass slide treatment: Microscope coverslips (catalogue no. CS22X22C, 22 x 22 mm and 0.13–0.17 mm thickness, Livingston International, Broadmeadows, Australia) were etched in the top right corner using a 0.5 mm thick diamond tip scriber (catalogue no. MS62107-ST, PST ProSci Tech, Kirwan, Australia) and carefully placed on a coverslip holder (catalogue no. C14784, Invitrogen Molecular Probes, Eugene, Oregon, USA) using a specialised pair of tweezers (catalogue no. 51-1625-0165, Dumont HP crossover tweezers N7 Stainless steel curved 0.17 x 0.10mm tip, Oxford Instruments, Abingdon, Oxfordshire, England). Etching on coverslips allowed easy identification of the treated side of the coverslip and curved tip tweezers enabled its smooth handling throughout the ExM procedure. A holder containing coverslips was submerged overnight in 1 N hydrochloric acid solution (catalogue no. H9892, Merck, Irvine, United Kingdom) and sequentially washed in milliQ and 100% ethanol (catalogue no. 493546, Merck, Saint Louis, Missouri, USA) for 1 min each. Coverslips were then air dried followed by coating with 300 µL of poly-L-Lysine solution and incubated at RT 60 min. Excess poly-L-Lysine solution was pipetted off from the treated coverslips and washed under a steady stream of distilled water. Coverslips were air dried and stored individually in 6-well plates (catalogue no. 3516, Costar®, Corning, Glendale, Arizona, USA) at RT for up to a month.
8. Anchoring agents: Glutaraldehyde solution (GA, catalogue no. 354400, Merck KGaA, Darmstadt, Germany) was diluted to 0.25% (v/v) in milliQ water and stored at −20°C until use. Acryloyl-X SE (6-((acryloyl)amino)hexanoic acid, succinimidyl ester) (AcX, catalogue no. A20770, ThermoFisher Scientific, Eugene, Oregon, USA), solution was prepared at 10 mg per mL concentration (w/v) in dimethyl sulfoxide (DMSO, catalogue no. 47230, Wuxi, Jiangsu, China) and stored in −20°C only up to six months.
9. Gel recipe 1: Polymer solution 1 was adapted from Wen et al. (2021) and contained 2.5% of sodium acrylate (SA; catalogue no. 408220, Merck, Milwaukee, Wisconsin, USA), 8.6% of acrylamide (AA), 0.15% of cross linker (N,N’-Methylene-bis-acrylamide, BIS; catalogue no. 146072, Merck, Milwaukee, Wisconsin, USA), 2 M of sodium chloride (NaCl, catalogue no. S9888, Saint Louis, Missouri, USA) prepared in 1X PBS. Gelation solution was freshly prepared prior to the gelation step by aliquoting 188 µL of polymer solution 1 with sequential addition of 2 µL of milliQ water, 2 µL of 4-Hydroxy-2,2,6,6-tetramethylpiperidine 1-oxyl (TEMPO, catalogue no. 176141, Merck, Buchs, Switzerland; final concentration 0.01%), 4 µL of ammonium persulfate (APS, catalogue no. 09913, Merck, Buchs, Switzerland; final concentration 0.15%) and 4 µL of N,N,N’,N’-tetramethylethane-1,2-diamine (TEMED, catalogue no. 17919, Thermo Fisher Scientific, Waltham, Massachusetts, USA final concentration 0.15%).
10. Gel recipe 2: Polymer solution 2 was adapted from Gambarotto et al. (2021) which contained 23% of sodium acrylate, 10 % of acrylamide, 0.1 % of crosslinker (BIS) prepared in 1X PBS. Gelation solution 2 was freshly prepared prior to the gelation step by sequentially mixing 180 µL of polymer solution 2 with 10 µL of TEMED and APS (final concentration 0.5 % each). Both polymer solutions 1 and 2 were made at least a day in advanced and stored at −20°C for no more than 2 weeks to avoid fragile gels.
11. Denaturation buffer: The denaturation buffer was comprised of 5% SDS, 50 mM of Tris-HCL, 1 mM of EDTA, 1 M of NaCl, 0.15 % Triton X-100 prepared in milliQ. Some trials involved addition of 8 M urea (substituting for SDS) in denaturation buffer while other components were added at above concentrations. This buffer was stored at room temperature (RT) and pre-warmed to 30°C prior to its use.
12. ExM wash buffer: The washing buffer was prepared by mixing 1% of Triton X-100 in 3X PBS buffer. This buffer was stored at RT until use.
13. Homogenisation buffer: The homogenisation buffer was made by mixing 50 mM of Tris-HCL (catalogue no. 648317, Merck, KGaA, Darmstadt, Germany) with 0.25%Triton^TM^ X-100, 0.8 M of guanidine (catalogue no. G3272, Merck, Saint Louis, Missouri, USA), and 2 mM of calcium chloride (CaCl_2_, catalogue no. 21101, Buchs, Switzerland) in 3X PBS. This buffer was stored in 4°C for up to two weeks.
14. Gel storage solution: Propyl gallate (PG, catalogue no. P3130, Milwaukee, Wisconsin, USA) powder (0.2% w/v) was added in milliQ water and dissolved by gentle mixing on a rotary mix and stored at RT only up to a month. This solution was discarded if turned yellow.
15. Antibody stocks: The primary calmodulin monoclonal antibody 6D4 (Catalogue no. MA3-918, Invitrogen, Carlsbad, California, USA) was diluted (1:500 from 1 mg per mL stock) in 3X PBS and stored in smaller aliquots of 10 µL per vial in −20°C. The secondary antibody, anti-mouse Alexa Fluor^TM^ 546 (catalogue no. A-11003, Invitrogen, Carlsbad, California, USA) was freshly diluted (1:50 from 1 mg per mL stock) in 3X PBS and mixed with 10% of Image-IT^TM^ FX signal enhancer prior to the post labelling step.
16. Non-specific fluorescent labels: For non-specific protein labelling, Alexa Fluor™ 488 NHS Ester (catalogue no. A20000, Life Technologies, Eugene, Oregon, USA) was diluted in DMSO to prepare 1 mg per mL of stock. Aliquots of 6 µL was stored in PCR vials to avoid repeated freeze thaw. For nucleic acid labelling, ready to use stocks of DAPI (10 mg per mL) and SYTOX^TM^ blue (1 mM) were commercial procured (catalogue nos. H3569 and S34857, Life Technologies, Eugene, Oregon, USA). These were stored as per manufacturers recommendation.
17. Conjugated fluorophores: DBCO-modified TAMRA (ex/em max: 548/562 nm; catalogue no. A131, Click Chemistry Tools, Beijing, China) and Azdye 594 (ex/em max: 590/617 nm; catalogue no. CLK-1298, Jena Biosciences, Deutschland, Germany) stocks were prepared at 30 mM concentration in DMSO and stored in −20°C until its use. Alexa Fluor^TM^ 488 conjugated to streptavidin molecule (ex/em max: 495/519 nm; catalogue no. S11223, Life Technologies, Eugene, Oregon, USA) was diluted in 1X PBS to prepare 2 mg per mL of working stock and aliquots of 10 µL were stored in −20°C until its use. Phalloidin-iFluor 488 conjugate (catalogue no. ab176753, Abcam, Cambridge, United Kingdom) was diluted (1:500) in 1.5 mL of 3X PBS prior to staining.
18. ExM reagents: The thermoaffinity tri-functional chemical linkers, 1 (ER-60-308) and 2 (ER-60-309) were supplied by Chrometra Scientific, Leuven, Belgium. These compounds (Figure S1) were separately diluted in DMSO to prepare 33 mM of working stock and stored at −20°C until its use.
19. Oligonucleotide probes: The modified and unmodified libraries (Table 1) of oligonucleotide probes targeting 18S rRNA (5′ GGGCAAGATCG AAAACTTG 3′) of *B. minutum* (Yokouchi et al., 2003) was procured from Integrated DNA Technologies, Coralville, Iowa, USA. The modified oligonucleotide libraries harboured Acrydite™ moieties on the 5’ end, which enables covalent grafting of the hybridised product in hydrogel, whereas its 3’ end contained biotin molecule that permitted post labelling of the product using a fluorophore conjugated with streptavidin. Similarly, a separate oligonucleotide library with primary amine modification on 5′ reactive with tetrafluorophenyl ester group of linker 2 was commercially obtained. All oligonucleotide libraries were diluted in DNAse free water to prepare 10 mM of working stock.

### Coupling of modified oligonucleotides with trifunctional linkers

The oligonucleotide libraries (details of modification in Table 1) were separately coupled with platinum (II)-based (linker 1) and tetrafluorophenyl ester group containing (linker 2) compounds as described in Wen et al. (2021). Briefly, an aliquot of 1.5 µL of unmodified FISH oligonucleotide probe (2.67 mM of working stock) was mixed with 1.2 µL of linker 1 (33 mM working stock) along with 90 µL of milliQ water and 10 µL of DMSO. Linker 1 compound was not coupled with any fluorescent dye beforehand.

**Table 1.** Commercially available oligonucleotide modifications, trifunctional linkers and modified fluorophores to integrate FISH with ExM for 18S rRNA labelling in Breviolum minutum.

| Oligonucleotide modification | Linkers** | Compatible fluorophores |
| --- | --- | --- |
| Unmodified | 1* | DBCO (e.g., Azdye 594 DBCO in this manuscript) |
| Primary amine on 5' | 2 | DBCO (e.g., TAMRA DBCO in this manuscript) |
| 5' Acrydite™ and 3' biotin* | No linker | Streptavidin conjugated with Alexa fluor 488™ |

\*indicates post-ExM labelling strategies; \*\*details of linkers are outlined in supplementary Figure S1.

Unlike linker 1, linker 2 compound and DBCO-modified TAMRA dye were separately diluted in DMSO to prepare 2 mM of working stock. Further, 200 µL of linker 2 compound was mixed with 240 µL DBCO-modified TAMRA dye and this mixture was incubated at RT on a shaker incubator (350 rpm) for 4 h. An aliquot of 200 µL of linker 2-DBCO-TAMRA was mixed with 44 µL of FISH oligonucleotide probe containing reactive primary amine (1 mM working stock concentration) in 5:1 ratio and incubated at RT for 2 h.

These mixtures were incubated at 55°C for 30 min on a shaker incubator (350 rpm). The oligonucleotide libraries functionalised with linker 2-DBCO-TAMRA and linker 1 were purified separately to remove any salts using Sephadex G-25 NAP 10 columns (catalogue no. 17085401, Global Life Sciences Solutions, Wilmington, Delawarem USA). For this, the total sample volume was adjusted to 1 mL using DNAse free water. Pre-existing storage buffer was passed through the columns by removing the top cap. Columns were equilibrated thrice using 5 mL of DNAse free water in each round. Filtrates were discarded and 1 mL of sample was carefully loaded on to the column. Samples were left to be completely absorbed in the column and then eluted using 100 µL of DNAse free water. The elution step was repeated 10 times and filtrate was collected in a fresh centrifuge tube for each round. The purity of each of these fractions was tested by recording their absorbance ratio at 260/280 nm. The fractions with absorbance ratio ranging from 1.75–1.85 were pooled and their yield was recorded (µg per mL). These yield values were used later to adjust the FISH probe concentrations (10 ng per µL) in the hybridisation step. The purified pooled fractions of oligonucleotides functionalised with the trifunctional linkers were stored at −20°C until use.

### Optimisation of ExM procedure

#### Sample fixation

A millilitre of exponentially growing cultures of *B. minutum* were harvested at 2 x 10^5^ cells per mL concentration and centrifuged at 10,000 *g* for 5 min. Supernatant was discarded, and cell the pellets were washed with fRSS. The washed cell pellets were separately treated with combinations of chemical fixative agents, 4% PFA or 0.7% FA/1% AA, or 1.4% FA/2% AA for 30 mins. The fixed samples were centrifuged at 10,000 *g* for 5 min and washed twice using 500 µL of fRSS. Supernatants were discarded, and cell pellets were resuspended in 100 µL of 3X PBS. These cell suspensions were sonicated in a water bath (Power Sonic 505, Thermoline Scientific, Wetherill Park, Australia) set at 40 Hz for 10 min. Samples were further centrifuged at 10,000 *g* for 5 min and cell pellets were used for pre-ExM labelling either using fluorescence *in situ* hybridisation (FISH) procedure or primary antibody labelling.

#### Pre-ExM labelling - Fluorescence in situ hybridisation and antibody

For FISH, cell pellets were resuspended in 80 µL of hybridisation buffer containing the 18S rRNA probe at 10 ng per µL concentration and incubated at 46°C for 3–4 h. Samples were centrifuged at 10,000 *g* for 2 min and supernatants were discarded. Cell pellets were resuspended in 500 µL of pre-warmed FISH wash buffer to remove any unbound probes. Samples were centrifuged at 10,000 *g* for 2 min and cell pellets were resuspended in 500 µL of pre-warmed FISH wash buffer. Samples were further incubated at 48°C for 20 min followed by centrifugation at 10,000 *g* for 2 min. Cell pellets were resuspended in 100 µL of 3X PBS. The labelled samples were treated with 500 µL of permeabilization buffer and incubated at RT for 60 min.

For antibody labelling, cell pellets were first resuspended in 500 µL of permeabilization buffer and incubated at RT for 60 min. Samples were then washed in 500 µL of 3X PBS and blocked at RT using 200 µL of Image-IT^TM^ FX signal enhancer (catalogue no. I36933, Life Technologies, Eugene, Oregon, USA). Samples were further centrifuged at 10,000 *g* for 2 min, supernatants were removed, and cell pellets were washed twice using 500 µL of 3X PBS. The primary antibody was added in samples at 1:500 dilution along with 10% of Image-IT^TM^ FX signal enhancer prepared in 3X-PBS. Samples were then incubated overnight at 37°C on a rotary shaker. Pre-ExM FISH and antibody labelled samples were washed twice using 500 µL of 3X PBS and centrifuged at 10,000 *g* for 2 min. Cell pellets were resuspended in 100 µL of 3X PBS and stored in −20°C until the next day.

#### Anchoring

Samples were thawed and treated separately with 10 µL of anchoring agents, AcX (0.01 mg per mL final concentration) or GA (0.025% final concentration) for 30 min at RT followed by centrifugation at 10,000 *g* for 2 min. Supernatant was discarded and cell pellets were washed twice using 500 µL of 3X PBS. Cells were finally suspended in 100 µL of 3X PBS and carefully deposited on acid-washed coverslips coated with poly-L-Lysine. Cells were allowed to adhere to treated coverslips for 60 mins at RT. At the end of this incubation step, excess solution was aspirated using a micro-pipette and discarded. No anchoring was performed for samples fixed with different percentages of FA/AA (0.7/1 or 1.4/2) because these chemicals are sufficient for the modification of biomacromolecules for their covalent grafting in hydrogels (Gambarotto et al., 2021).

#### Gelation

Coverslips coated with cells were separately pre-infused with 100 µL of ice-cold polymer solutions 1 or 2 and further incubated on an ice bath for 20 min. Gelation solutions 1 and 2 were prepared freshly in separate centrifuge tubes that were pre-chilled on a benchtop cooler box (catalogue no. 355501, Thermo Scientific^TM^, New York, USA) stored in −20°C overnight. The coverslips coated with cells overlaid with polymer solutions were gently lifted using Dumont curved tweezers and slightly tilted to remove any excess solution. These coverslips were placed on the ice bath and again covered separately with of ice-cold gelation solution 1 or 2. An aliquot of respective gelation solutions 1 or 2 (90 µL) was carefully added as a drop in-between two spacer coverslips (22 x 32 mm, 1.5 mm thickness; catalogue no. CS22X32-1.5, Livingston International, Broadmeadows, Australia) placed on the either ends of Sigmacoat^TM^ treated glass slide. The main coverslip was then placed (upside down) on the drop of the gelation solution ensuring that the surface of the coverslip coated with cells (i.e., etched surface) was in contact with the solution. Four small binder clips (15 mm) were used to firmly hold down the main coverslip with two spacer coverslips placed over a glass slide (step 4 in Figure 1). This assembly was then placed in a humidified chamber (airtight box lined with wet tissue paper) and incubated at 37°C for 2 h. Azpack^TM^ carbon razor blades (Fisher Scientific, catalogue no.11904325, Leicestershire, United Kingdom) were carefully inserted between the spacer coverslips and glass slide on the either ends and slowly moved towards the centre to separate coverslips in order to expose the thin film of gel formed on one side of the coverslips. This continuous piece of gel covering the main coverslip (in middle) and spacers (on sides) was carefully cut to remove the spacer coverslips. The gel formed on the centre of the main coverslip was roughly cut into square shape of ∼ 0.5 x 0.5 mm dimension and its top right corner was trimmed. This helped in maintaining the correct orientation and placement of the correct side of the gel (cells in close contact with imaging chamber) for post-ExM imaging. The gel piece was carefully separated from the main coverslip using a thin plastic sheet (0.1 mm thickness).

**Figure 1.**
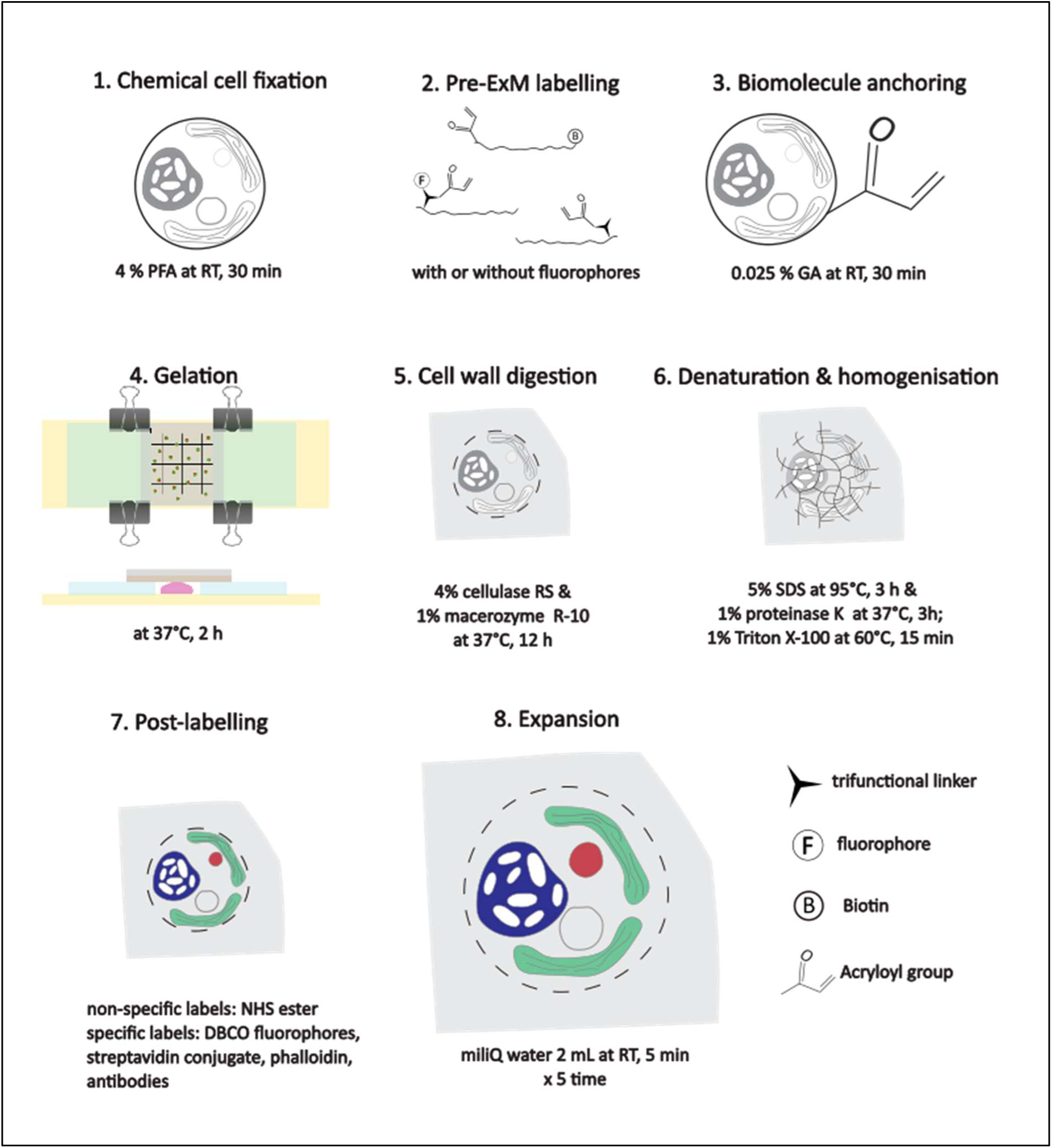
Overview of the optimised expansion microscopy workflow for 4-fold expansion of the cnidarian photosymbiont, *Breviolum minutum*.

#### Cell wall digestion

The gel piece was carefully transferred from the plastic sheet using a paint brush (size 0) to a well containing 2 mL of cell wall digestion enzyme mix in a 6-well plate. Gels were further incubated at 37°C on a shaking incubator (50 rpm) for 2, 4, and 12 h. The efficacy of cell wall digestion for cells suspended in liquid (PBS) prior to the anchoring step was also tested separately as part of the optimisation. For this, fixed cell samples were centrifuged (10,000 *g* for 2 min) and washed twice using 3X PBS. Supernatant was discarded and the cell pellet was directly resuspended in 1 mL of cell wall enzyme digestion mix supplemented with 0.5 M D-sorbitol. Samples were incubated in the dark at 37°C for 6, 12 and 24 h. Then samples were centrifuged (200 *g* for 2 min) and the cell pellet was washed in regeneration buffer (0.5 M sucrose, 0.5 M sorbitol, 25 mM of CaCl_2_ prepared in 3X PBS). Samples were incubated in the dark for 3 h at RT and centrifuged (200 g for 2 min) to obtain a cell pellet which was washed thrice in 500 µL of 3X PBS.

#### Thermo-chemical denaturation

The next day, the enzymatic mix was removed with a pipet, and gels were washed twice using 1 mL of 3X PBS for 5 min. Denaturation buffer was pre-warmed at 30°C and 2 mL were dispensed in each well. Gels were separately denatured for 1.5, 3, and 12 h at 37°C and 55°C, and for 1.5, 3 h at 95°C. The latter condition was not chosen for 12 h incubation due to excessive drying of the thin gels. A total of eight different combinations of temperature, time and denaturation chemical were tested to achieve isotropically expanded spherical *B. minutum* cells. The denaturation step was terminated by replacing denaturation buffer with 2 mL of ExM wash buffer and incubated at 65°C for 20 min. Gels were further washed twice using 2 mL of ExM wash buffer and incubated at RT for 5 min. This washing step is important to remove SDS which might denature the enzyme (proteinase K) added in the next homogenisation step. Denaturation was also tested separately using 8M urea at 37°C incubated for 4, 12 and 48 h.

#### Homogenisation

The chemically denatured gels (SDS and urea) were further digested in homogenisation buffer containing 80 U of proteinase K enzyme (catalogue no. P8107S, New England Biolab, Ipswich, Massachusetts, USA) and separately incubated either for 1.5 and 3 h at 37°C or 58°C. We also separately tested overnight homogenisation using either SDS or proteinase K at RT, 37 and 58°C without any prior denaturation (Figure S7). The homogenisation step was terminated by washing of gels thrice in 1 mL of 3X PBS and incubated for 5 min at RT. Gels were stored in 4°C overnight in 1.5 mL of 3X PBS in case fluorescence labelling was not possible to be performed on the same day.

#### Post-labelling and Expansion

The next day gels were brought back to RT and its pre-ExM dimensions were measured using a Vernier calliper. A reactive dye, Alexa Fluor™ 488 NHS Ester was added at 4 µg per mL concentration for non-specific staining of proteins. Gels containing FISH labelled *B. minutum* cells with linker 1 trifunctional linker were detected separately using 5 µM of DBCO-modified Az594 dye diluted in 2 mL of 3X PBS, and incubated in dark at 26°C, 50 rpm for 1 h. FISH labelled samples containing biotin modification on 3′ end of the FISH probe were detected using 2 µg per mL of streptavidin conjugated with Alexa Fluor^TM^ 488 dye prepared in 2 mL of 3X PBS, and incubated in dark at 26°C, 50 rpm for 1 h. Gels containing samples with primary antibody were separately stained in 1.5 mL of secondary antibody solution (1:50 dilution) and incubated in dark at 37°C, 50 rpm for 2.5 h. These gels were washed thrice using 0.1% Tween 80 dissolved in 3X PBS. Some gels were post-labelled separately using 1.5 mL solution of Phalloidin-iFluor 488 conjugate and incubated at 26°C, 50 rpm for 2.5 h.

Gels were expanded five times by incubation in 2 mL of milliQ water for 10 min in each round. The dimensions of the expanded gels were measured using a Vernier calliper to determine the expansion factor of the gel. In case imaging of the expanded gels was not possible on the same day, they were stored in 2 mL of PG solution at 4°C in dark conditions.

In few samples, the efficacy of cell wall digestion pre- and post-ExM was assessed using 20 µg per mL of calcofluor white (catalogue no. 18909, Merck, Buchs, Switzerland) prepared in milliQ water. Some gels were also subsequently labelled with nucleic acid stains, 15 µg per mL of DAPI or 2 µg per mL of SYTOX^TM^ Blue in milliQ water for 15 min at RT. Excess unbound stains were washed twice in 2 mL of milliQ water (3X PBS in case of pre-ExM labelled samples).

### Pre-validation of fluorophore labelling in *B. minutum*

All specific (trifunctional linkers coupled with modified oligonucleotide for FISH, calmodulin antibody, and phalloidin conjugate) and non-specific (NHS ester) fluorescence labelling strategies were pre-validated using FLIM in *B. minutum* as described in (Deore et al., 2024a, 2022) prior to ExM optimisation. Briefly, we utilised unique positioning of autofluorescence signals on the phasor plot (a two-dimensional representation of phase vectors G and S) indicative of its lifetime which is distinct from exogenous fluorophores (for pre-ExM validation). We used pure exogenous fluorophores (TAMRA, Az594, Alexa Fluor^TM^ 488, iFluor 488, and Alexa Fluor^TM^ 564) wherever possible to ensure that signals arise from these were uniquely located on phasor plot and are distinct from autofluorescence signature. We used pure bacterial cultures to validate FISH labelling using 16S rRNA oligonucleotide (EUBmix 338: 5′ GCTGCCTCCCGTAGGAGT 3′) coupled with trifunctional linkers and fluorophores (linkers 1 and 2 with Az594 dye) as well as oligonucletides with biotin modification (detected using Alexa Fluor 488 conjugated to streptavidin). We further used these FISH labelled bacterial cultures to obtain unique signatures on phasor plot for Az594, streptavidin conjugated Alexa Fluor 488 signatures using FLIM.

### Imaging

A specialised polymer coverslip bottom µ-Slide 2 Well slides (#1.5, catalogue no. 80286, ibidi, Planegg, Germany) were coated with poly-L-Lysine solution at RT for 30 min. The excess solution was removed by pipetting, and the surface was rinsed once with milliQ water. The gels were carefully placed on to the treated µ-Slide bottom while ensuring that the correct surface of the gel containing *B. minutum* cells was in contact with the imaging surface (cut right top alignment). A glass coverslip (18 x 18 mm) was added onto the top of the gel to completely flatten it against the surface and ∼100 µL of milliQ water was added surrounding the gel to avoid drying. The mounted gels were imaged on a Leica TCS SP8 Lightning confocal system running LAS X using a 20x HC PL APO NA 0.75 dry objective. Images were captured in 512 x 512-pixel resolution at 16-bit depth. Sample labelled using DAPI or SYTOX^TM^ blue stains were excited using 405 nm laser whereas Alexa Fluor^TM^ 488, iFluor 488 conjugates and Azdye 594, Alexa Fluor^TM^ 594 were excited using 488 and 564 nm lasers, respectively. Detectors were set to emission ranges 450–500 nm, 510–550 nm, 600–650 nm.

### Image and statistical analysis

All raw images were processed in Fiji (v1.54s), an open-source image analysis platform based on ImageJ developed by National Institutes of Health, USA (Schindelin et al., 2012). Individual cell masks were created by manually drawing the regions of interests (ROIs) along the outer edge of a cell. These masks were used for the estimation of cell diameter and area. This information was used to calculate the perimeter and circularity (4π x area/perimeter^2^ where 1 corresponds to perfect circle and 0 is elongated shape) of pre- and post-ExM cells in R studio (2026.01.0 Poist software PBC). A separate line ROI was drawn over each cell to map the intensity of fluorescence signals (arbitrary units) across different position (microns) in pre- and post-ExM samples which is represented as line profile graphs in Figure 9. The raw data was inspected for assumption of equal variance and normality (Shapiro test) R studio using tidyverse, janitor, purr, dplyr, ggbeeswarm, and ggplot2 plugins. Welch two sample t-test was implemented to estimate the differences in mean cell diameter and circularity index between pre and pos-ExM cells. The level of significance was 0.05. The expansion factor of cells was estimated by dividing the mean cell diameter of post-ExM by pre-ExM cells.

## Results

### Sonication improves access to cellulosic thecal plates

The sonicated samples showed a better uptake of the cell wall stain, calcofluor white (blue coloured), revealing a thick and continous ring of cellulose rich material along the outer edges (amphiesma) of *B. minutum* cells compared to unsonicated control samples (Figure 2). While the cell wall staining improved in sonicated samples, the autofluoresence (red colour, Figure 2) diminished partly compared to control.

**Figure 2.**
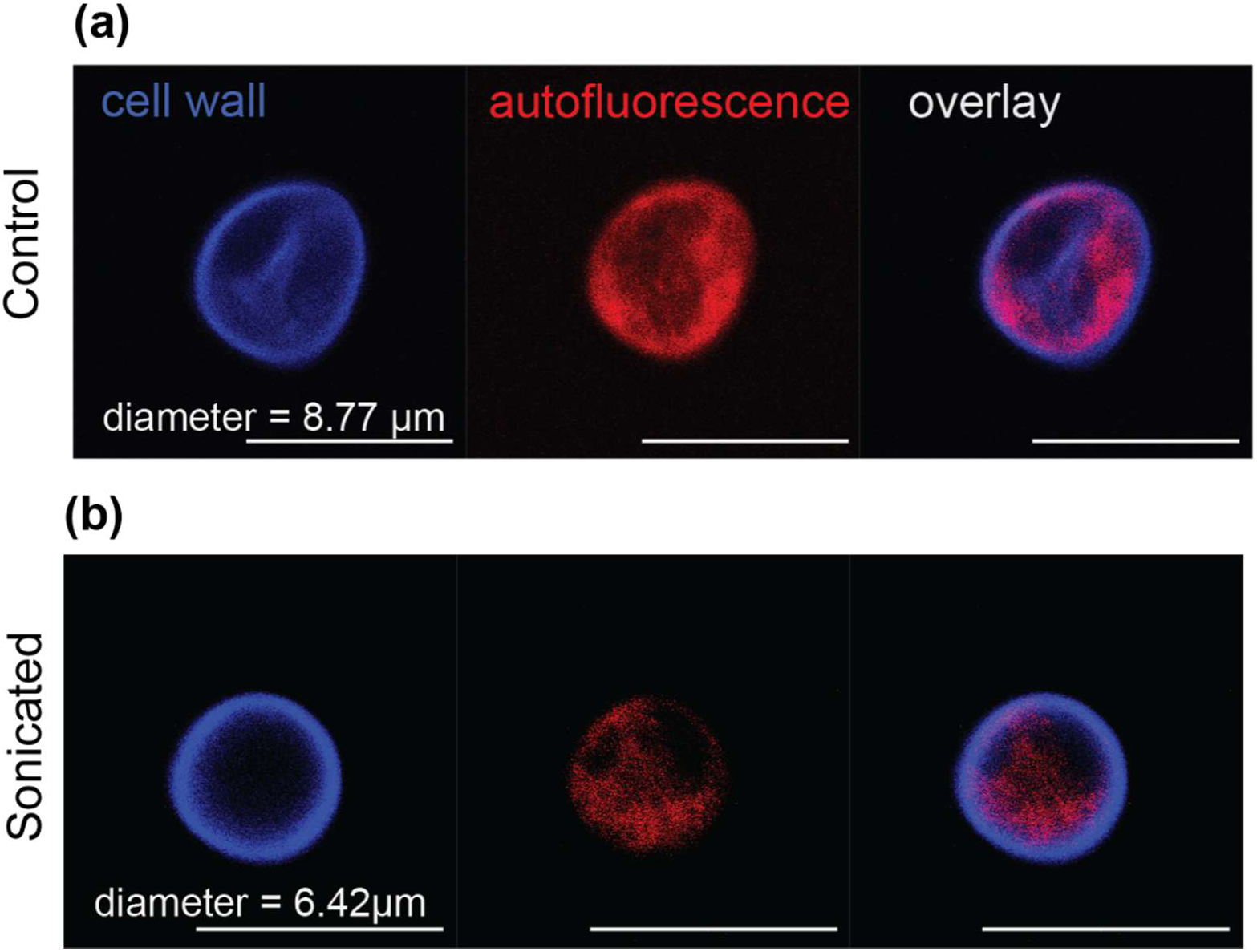
Pre-treatment of *B. minutum* cells in suspension. a) Control samples without sonication compared to b) cells sonicated for 10 min at 40 Hz. Blue coloured signals are from cell wall stained with calcofluor white and red represents autofluorescence. Overlay images are superimposition of blue and red channels. Scale bar = 10 µm.

### Protein dense ring surrounding pyrenoid in *B. minutum*

The separate chemical treatments for cell fixation and biomacromolecule anchoring using PFA and GA, respectively, resulted in homogeneous and consistent non-specific labelling of proteins (NHS ester) throughout the cell interior of *B. minutum* cells (green coloured signal in Figure 3a) compared to the samples treated with different percentages of FA/AA (0.7/1 or 1.4/2) which is a single step approach to achieving cell fixation as well as anchoring (Figures 3b and c, respectively). The NHS ester labelling approach combined with PFA fixation consistently revealed a protein dense ring structure which is hypothesised to be the pyrenoid, a carbon fixation organelle, of *B. minutum* (white arrow in Figure 3a). This feature was obscured in FA/AA treated samples (Figures 3b and c).

**Figure 3.**
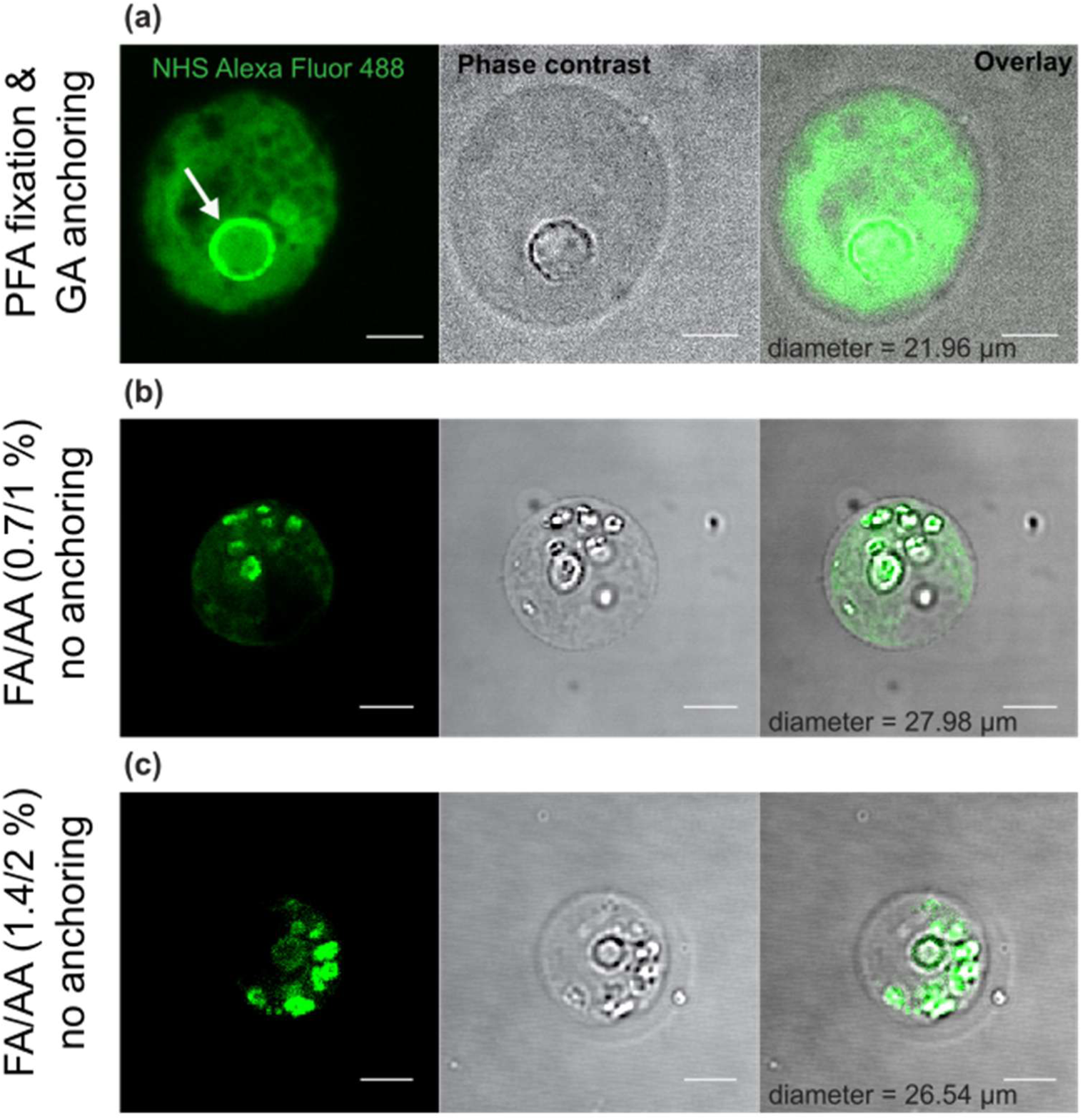
Post ExM images of *B. minutum* cells treated with different chemical fixatives (a) paraformaldehyde (PFA) fixed samples treated with glutaraldehyde-based (GA) anchoring in a separate step, (b) formaldehyde/acrylamide (FA/AA, 0.7/0.1 %), and (c) FA/AA (1.4/0.7 %) fixation without any separate anchoring treatment. Green coloured signals indicate non-specifically labelled proteins using NHS ester coupled with Alexa FluorTM 488. White arrow indicates protein dense organelle, pyrenoid. Overlay images are superimposition of green channel over phase contrast images. Scale bar = 10 µm.

### Glutraldehyde (GA) is a superior anchoring agent compared to acryloyl-X (AcX)

We compared two most commonly used biomacromolecule anchoring agents, GA and AcX, which facilliate embedding of cellular components within a hydrogel. samples treated with GA (Figures 4a and b) showed an overall better retention of the nucleus (blue coloured) and protein (green coloured) signals, and a circularity index of 0.8. Conversely, samples treated with AcX showed a loss of nuclei in gels separately stained with two dyes, DAPI and SYTOX^TM^ blue (Figures 4c and d). AcX treated samples also showed a substantial background interference outside of *B. minutum* cells. Overall, GA preseved a better cellular context with minimal background interference comapred to AcX treatment which only appears to be benificial in maintaing the circularity index of a cell (0.9).

**Figure 4.**
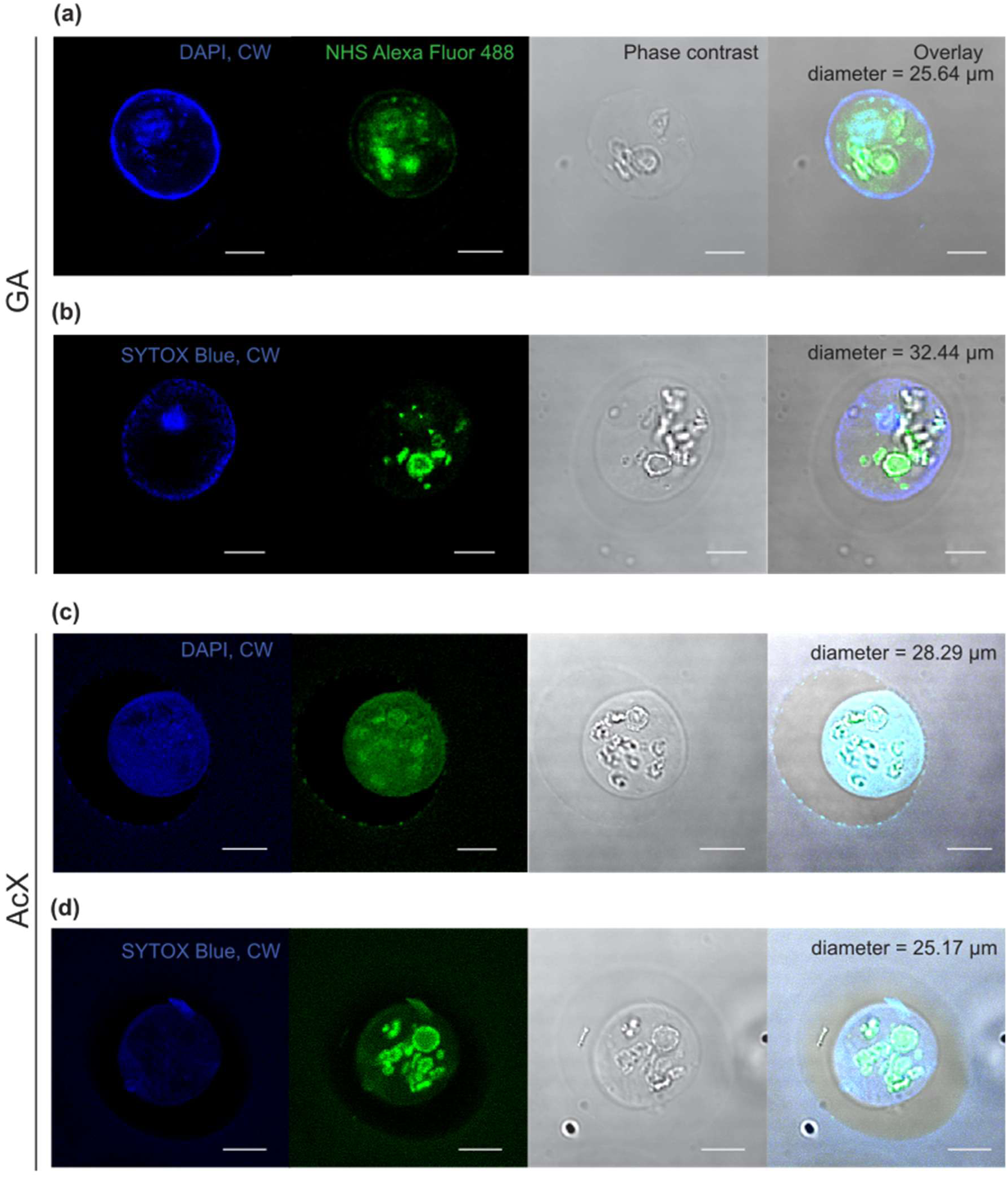
Post-ExM images of *B. minutum* cells treated with different anchoring agents, Glutaraldehyde (GA) and Acryloyl-X (AcX). Nucleic acid and cell wall are stained post-expansion using DAPI or SYTOXTM Blue and calcofluor white (CW) dyes, respectively (blue coloured signal). Green coloured signals are non-specifically labelled proteins using NHS ester coupled with Alexa FlourTM 488. Overlay images are superimposition of blue and green channel over phase contrast images. Scale bar = 10 µm.

### Monomer to anionic polyelectrolyte molar ratio is crucial to improve expansion factor

The hydrogels prepared using gel recipe 1 contained higher percentages of anionic monomer (sodium acrylate, SA), relative to the neutral monomer (acrylamide, AA) compared to the gel recipe 2. The former recipe yielded ∼4.4-fold expansion of *B. minutum* cells (n=13, Table S1) and better retention of a cellular organelle, pyrenoid (white coloured arrow, Figure 5a). Whereas, gel recipe 2 resulted in only ∼3-fold expansion (n=6, Table S1) of cells and protein labelling showed distorted ring-like structures (green coloured, Figure 5a). The overall circularity index of post-ExM was 0.9 in samples prepared with either of the recipes and showed not significant difference comapred to pre-ExM cells.

**Figure 5.**
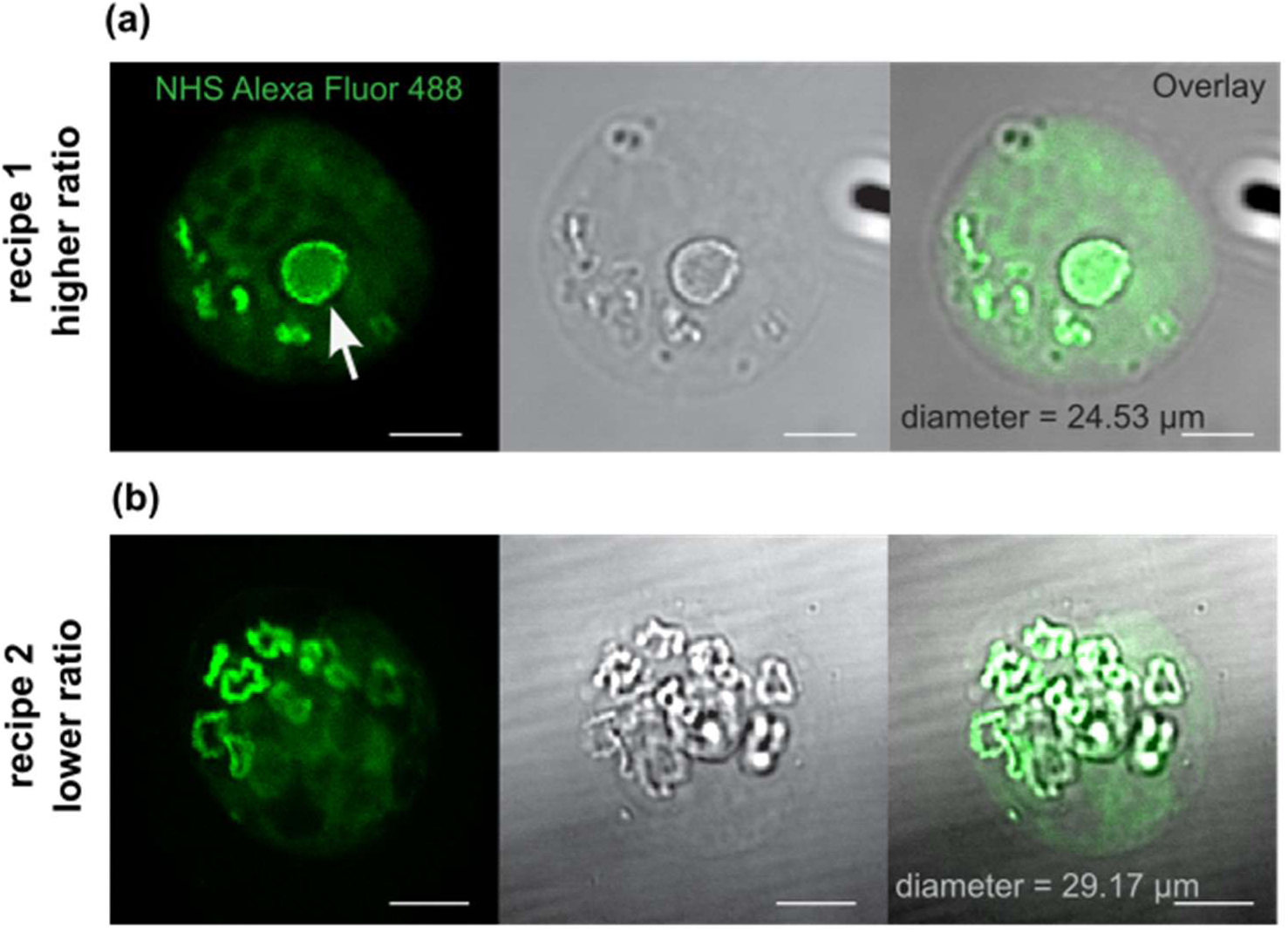
Optimisation of ExM gel recipe through varying monomer (acrylamide) to anionic polyelectrolyte agent (sodium acrylate) ratio to achieve isotropic expansion of *B. minutum* cells. a) gel recipe 1 (adapted from Wen et al. 2021) and (b) gel recipe 2 (adapted from Gambarotto et al. 2021). White arrow indicates a well preserved pyrenoid structure in *B. minutum*. Overlay images are superimposition of green channel over phase contrast images. Scale bar = 10 µm.

### Efficient enzymatic cell wall distruption for cells embded in hydrogels

We tested for an enzymatic digestion of *B. minutum* cell wall in liquid using a mixture of cellulase and macerozyme to generate a protoplast as a starting material for ExM (before anchoring step). A strong calcofluor white signal indicating persistent cell wall (blue) was observed along with reduced autofluoresnce signals (red colour across across all samples incubated for 4, 6 and 12 h (Figure 6a). Post gelation (cells embeded in hydrogel) enzymatic digestion (Figure 6b) clearly showed partial cell wall digestion (discontinous blue signals) in samples incubated for 2 and 4 h. The prolonged incuabtion for 12 h resulted in comeplete digestion of cell walls which improved expansion factor by ∼4-fold (4 h vs.12 h samples in Figure 6b). We consistently observed that the omission of sonication combined with enzymatic cell wall digestion resulted in the complete lack of expansion of *B. minutum* cells.

**Figure 6.**
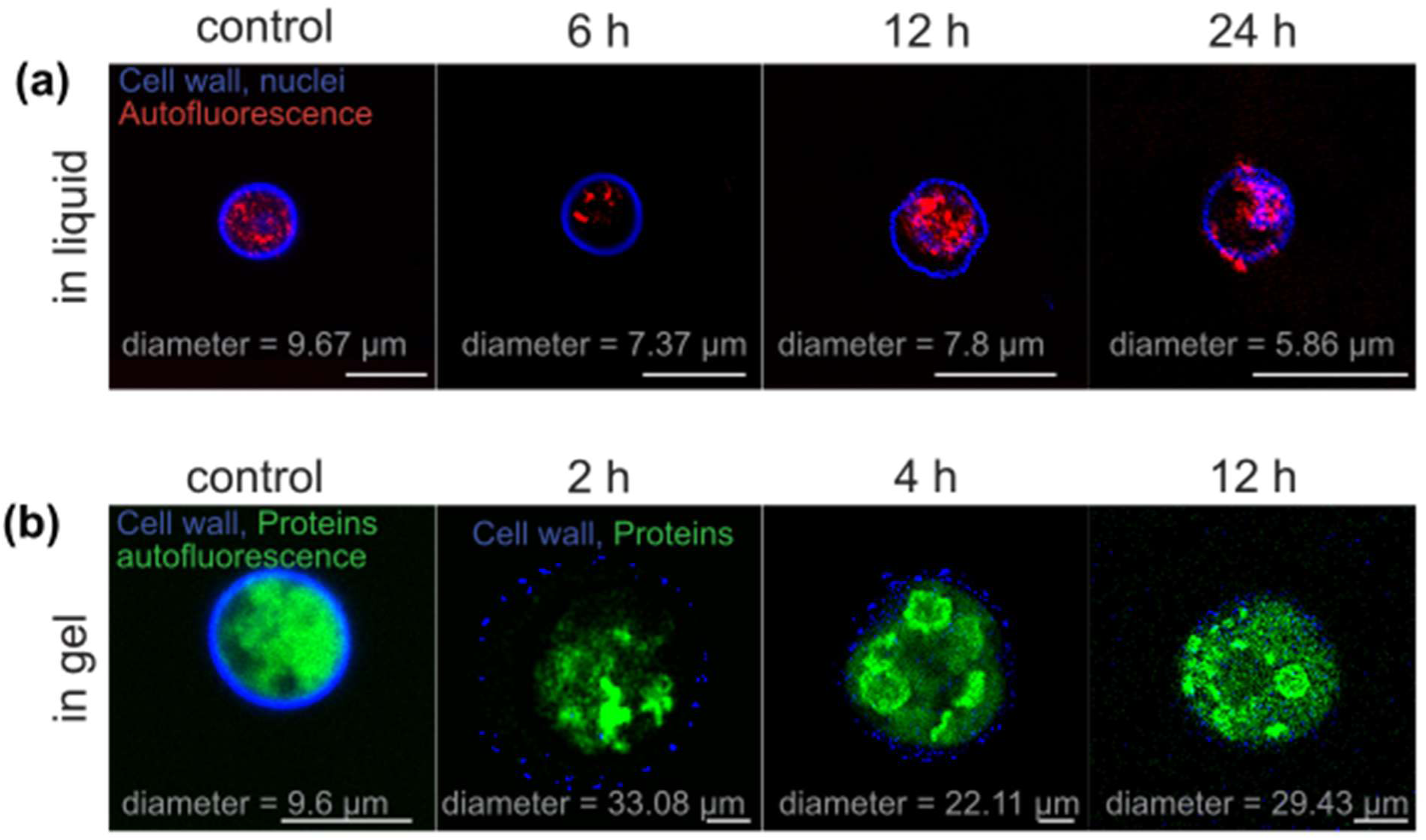
Enzymatic digestion of cellulosic cell wall in *B. minutum* cells. a) Unexpanded cells pre-treated with cellulase and macerozyme in liquid b) compared to *B. minutum* cells digested in hydrogels. All images depicted in panel a (in liquid) and control sample panel b (in gel) show unexpanded *B. minutum* cells. Only cell wall digested cells are expanded because of the potential steric hindrance exerted by cell walls. Blue coloured signals are from cell walls stained with calcofluor white (panels a and b) and DAPI (panel a) for nuclei labelling. Green signals are from non-specific labelling of proteins using NHS ester conjugated to Alexa Fluor 488 dye. Green signals in control *B. minutum* cell (in panel b) also contains autofluorescence signals which is indicated by red signals in panel a. Images are superimposition of blue and red or blue and green channels. Scale bar = 10 µm.

### Thermo-chemical denaturation combined with classical proteinase K-mediated homogenisation is key for 4-fold expansion in *B. minutum*

We tested a range of thermal conditions (37, 55 and 95°C), and chemical detergents (SDS and urea) for the denaturation of cellular components in *B. minutum*. This step was followed by classical proteinase K digestion treatment to enable isotropic expansion. In SDS treated samples (Figure 7a), we observed a moderate expansion of ∼2–2.5-fold in *B. minutum* cells at 37°C incubated for 1.5, 3 and 12 h however, this expansion appeared to be relatively non-isotropic as indicated by a circularity index lower than 0.8. The increase in denaturation temperature to 55°C led to 3-fold expansion and a circularity index of 0.9 in samples incubated for 3 and 12 h (Figure 7b). The prolonged incubation of 12 h at 55°C appears to severly impact the integrity of subcellular features. A consistent 3.5–4-fold expansion of *B. minutum* cells was achieved at a denaturation temperature of 95°C and incubation time of 3 h (Figure 7c). The key subcellular feature, i.e., the pyrenoid, was well preserved in *B. minutum* cells incubated in these conditions. In samples treated with urea combined with a denaturation temperature of 37°C for variable incubation duration (4, 12, 48 h) we merely observed 2-fold expansion (Figure 7d).

**Figure 7.**
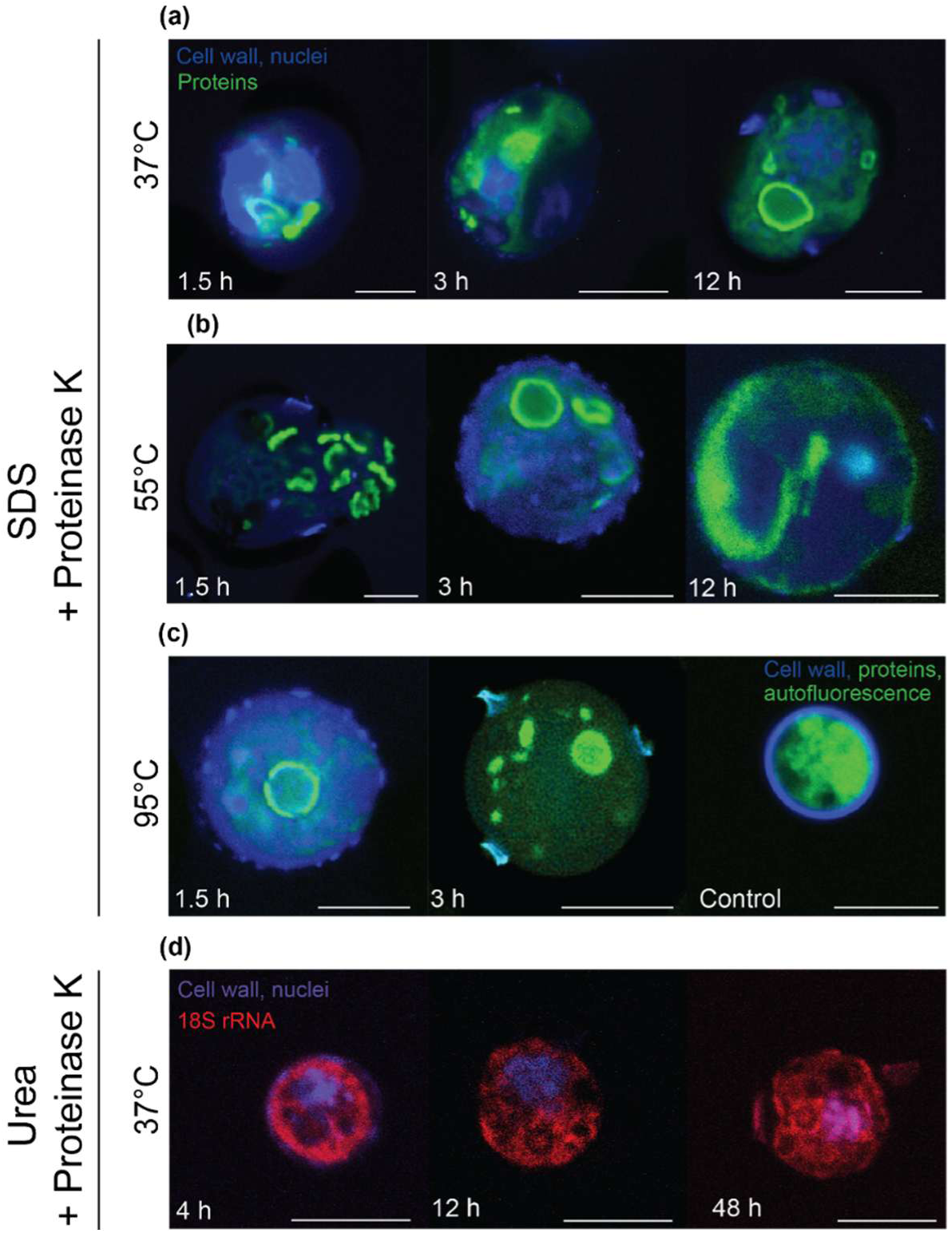
Optimisation of two step approach involving thermo-chemical denaturation conditions combined with proteinase K-mediated homogenisation (at 37°C for 3 h) for expansion of *B. minutum* cells. Detergent-based denaturation (a) using 5% Sodium dodecyl sulphate (SDS) incubated at 37°C (for 1.5, 3 and 12 h), (b) 55°C (for 1.5, 3 and 12 h), (c) 95°C (for 1.5 and 3 h), and (d) 8 M urea incubated at 37°C (for 4, 12 and 48 h). Green signals are from non-specific labelling of proteins using NHS ester conjugated to Alexa Fluor 488 dye. Blue signals are from cell wall stained with calcofluor white and DAPI for nuclei labelling. Red signal indicates (in panel d) FISH labelled 18S rRNA using C47 linker coupled with TAMRA. The control (in panel c) sample indicate unexpanded *B. minutum* cell wherein green signals indicate autofluorescence and non-specifically labelled proteins. Images are superimposition of blue and green (panels a-c), and red and blue (panel d) channels. Scale bar = 10 µm.

### A customised ExM approach to achieve 4-fold isotropic expansion of *B. minutum*

We tested a series of thermo-chemical and enzymatic combinations along with the duration of incubations to carefully resolve challenges associated with isotropic expansion of *B. minutum* cells (Figures 8a and b). We found that single step approaches that use either classical proteinase K-mediated homogenisation or detergent-based denaturation were ineffective and only led to 2.4 (n=3) and 2.7-fold (n=23) expansion, respectively. However, a combination of these two approaches which firstly involved detergent-based denaturation (5% SDS) at 55°C for 1.5 h followed by proteinase-K mediated homogenisation at 37°C for 1 h helped increase the expansion factor to 3-fold (n=13). Further increasing the duration incubations both for denaturation and homogenisation steps upto 3 h helped to achieve 3.5-fold expansion factor (n=23) while maintaining the circularity index closer to 0.9 (Figure 8, Tables S1 and S2).

**Figure 8.**
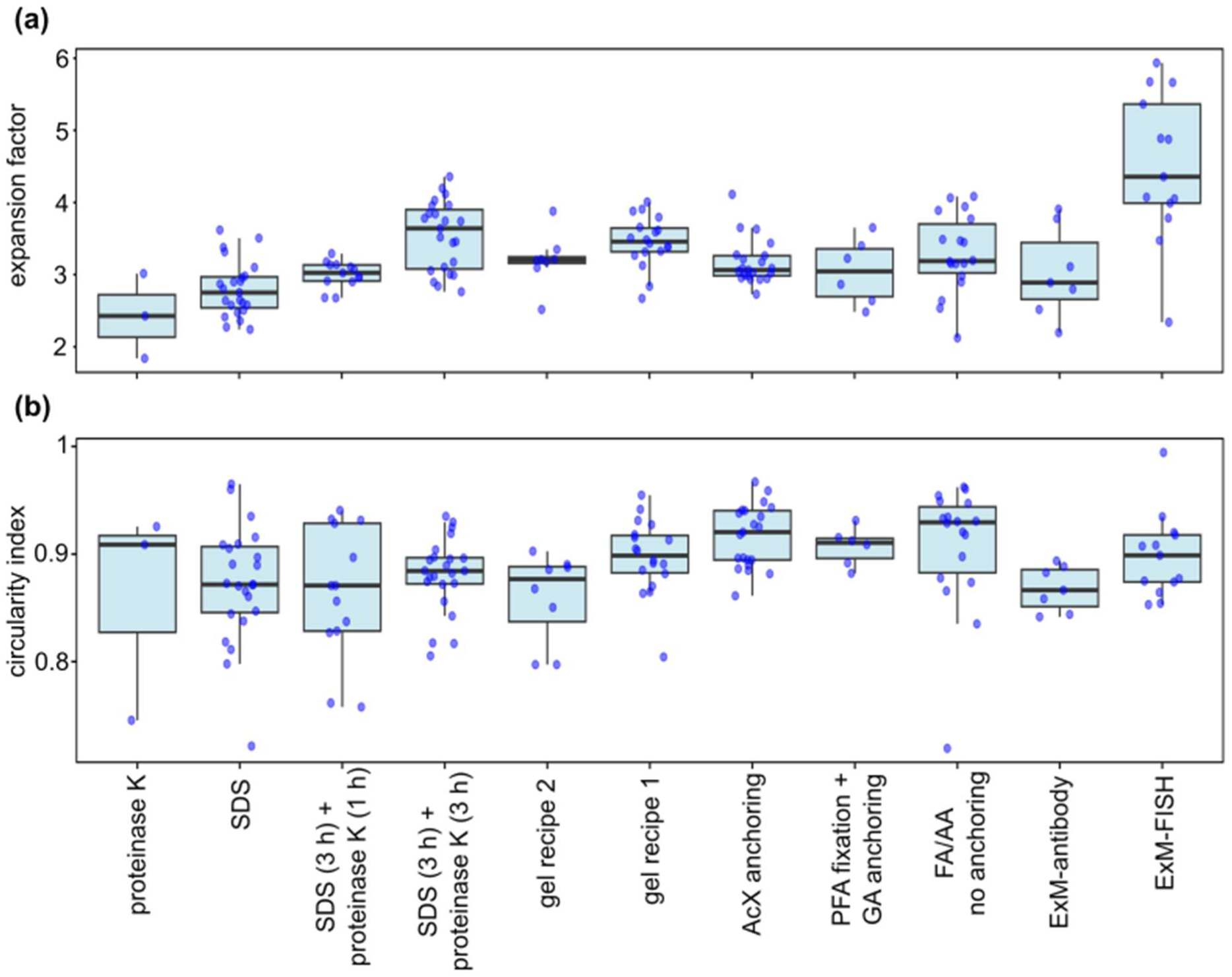
Customisation of thermo-chemical and enzymatic conditions for the improvement of (a) expansion factor while maintaining (b) the circularity index (i.e., isotropy) of *B. minutum* cells. Box and whisker plots show data median as horizontal central bar whereas top and bottom lines are upper and lower quartiles, respectively blue dots indicate individual cells and vertical line show dispersion.

Further, we explored two different gel recipes contaning different molar ratios of neutral monomer (AA) to anionic monomer (SA). The gel recipe 2 with lower SA concentrations relative to AA (Gambarotto et al., 2021) led to only 3.2-fold expansion (n=8) as compared to the gel recipe 1 (Wen et al., 2021) which resulted in 3.5-fold expansion (n=18). We also found that the most commonly used anchoring agent, AcX only yielded 3.1-fold expansion (n=21) although with better preservation of circularity index (0.9) compared to GA.

We tested two approaches for cell fixation to preserve the cellular ultrastructure. Our first approach involved PFA treatment prior to the ExM procedure (Laporte et al., 2022), while the second approach combined both cell fixation as well as biomolecule anchoring through FA/AA treatments (Gambarotto et al., 2021). Both of these approaches yielded 3-fold expansion with circularity index >0.9.

Based on above observations we narrowed down a strategy combining PFA-based cell fixation, gel recipe 1, and a two step thermo-chemical denaturation followed by classical proteinase K-mediated homogenisation for *B. minutum* exapansion. We implementated this approach to test compatibility of this protocol for post-ExM antibody labelling for the detection of calmodulin proteins, and visualisation of the 18S rRNA pool (FISH labelling) using tri-functional linker 1. We observed that antibody labelled cells were only 3-fold expanded with circularity index of 0.86 whereas, FISH labelled samples were expanded by 4.4-fold with circularity index of 0.9.

### Trifunational linkers and click-chemistry labelling enables integration of FISH with ExM

We used FISH labelling approach to label the 18S rRNA pool in the cytoplasm of *B. minutum* cells. These hybridised products were covalently bound to the hydrogel through acryloyl reactive groups, mimicking the monomer in trifuntional linkes 2 and 1 (Wen et al., 2021).The former linker contained the DBCO-modified fluorophore, TAMRA attached to 18S rRNA hybridised product in *B. minutum* cells prior to ExM procedure. Whereas, 1 linker attached to the 18S rRNA target contained a reactive azide group which was exploited to attach a DBCO-modified Az594 fluorophore post-ExM procedure. The FISH labelled samples (Figures 9a and b) contaning both linkers showed strong signals in pre-ExM samples. However, the samples containing linker 2 coupled with TAMRA signals showed a substantially subdued line profile indicating dampened fluorescence siganls in post-ExM samples (Figure 9a). The post-ExM line profiles of samples containing linker 1 coupled with Az594 fluorophore showed spatially well resolved and subatatially higher signals compared to pre-ExM *B. minutum* cells (Figure 9b). The non-specific protein labelling using NHS ester coupled with Alexa Fluor^TM^ 488 showed a consistent and homogeneous labelling within cytoplasmic spaces (Figure 9c) which is distinct from the 18S rRNA pool (Figures a and b) of *B. minutum*.

**Figure 9.**
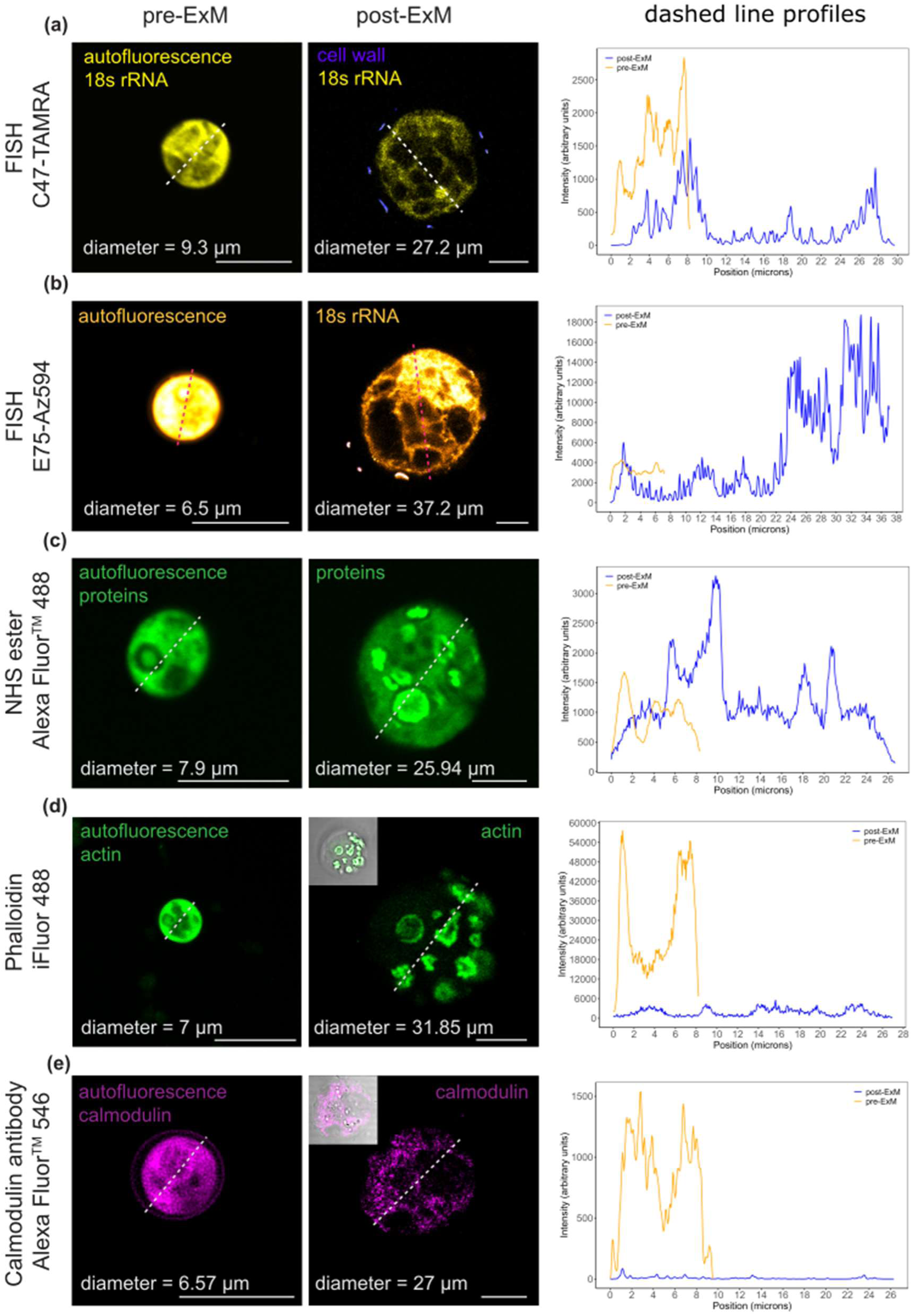
Pre and post-ExM labelling strategies for the visualisation of subcellular features in *B. minutum*. (a) Pre-ExM FISH labelling of 18S rRNA using C47 linker (tetrafluorophenyl ester) coupled with DBCO modified TAMRA. (b) Post-ExM fluorescence labelling of oligonucleotide hybridised with 18S rRNA using E75 linker (platinum (II)-based) detected using DBCO modified Az594 dye, (c) non-specific labelling of proteins using NHS ester of Alexa Fluor 488, (d) labelling of F-actin (a cytoskeletal feature) using phalloidin conjugate, and e) monoclonal antibody labelling of calmodulin (calcium binding protein) in *B. minutum* cells. Images listed under pre-ExM panel are control cells in unexpanded state showing respective fluroescence labelling (except for b) along with autofluorescence. All images under post-ExM panels are expanded cells devoid of any autofluorescence and only contains respective fluorescence labels added pre (in a) or post-ExM (b–e). Line profiles correspond to pre- and post-ExM fluorescence signal (orange and blue-coloured lines) positioned along the dotted lines overlayed on cells presented in adjacent images. The inset image in panel e (post-ExM) is an overlay image showing the distribution of calmodulin proteins with the context of *B. minutum* cell in phase contrast image. Scale bar = 10 µm.

We also observed strong fluorescence signals that were distinct from autofluorescence in *B. minutum* cells FISH labelled using biotin modified 18S rRNA oligonucleotides and detected by streptavidin conjugated to Alexa Fluor 488. A few cells in the negative control containing unlabelled *B. minutum* cells and streptavidin conjugate (no FISH probe) also showed fluorescence signals (supplementary Figure S3c). We observed a similar patterns in pure bacterial cultures FISH labelled using 16S rRNA and unlabelled samples containing only streptavidin conjugate with the fluorophore (supplementary Figures S3d and e).

### Visualisation of actin patches and calmodulin proteins in *B. minutum*

We used a phalloidin molecule coupled with iFluor 488 for the visualisation of the F-actin cytoskeleton and a monoclonal antibody to detect calmodulin proteins (calcium binding) in cytoplasmic spaces of *B. minutum*. The phalliodin labelling revealed F-actin patches (Figure 9d) as opposed to our expectations of visualising cytoskeletal filaments. The overall apperance of these F-actin patches is strikingly similar to the cytoplasmic ring-like structures observed in cells treated with non-specific protein label NHS ester (Figure 9c). A weak phalloidin signal was also observed in close proximity of the protein-rich ring surrounding the pyrenoid in *B. minutum* (Figure 9d). The antibody labelling revealed a distinct signal indicating the presence of calmodulin protein in the plasma membrane of pre-ExM *B. minutum* cell (Figure 9e). However, in post-ExM cells, calmodulin signals were heterogenously distributed in cytoplasmic spaces. The line profiles in Figures 9d and e showed a substantially lower, but well resolved, signal in post-ExM samples.

## Discussion

*B. minutum* is a widely studied microbial symbiosis model system for understanding its association with bacteria (Maire et al., 2021) and cnidarian hosts (Nitschke et al., 2022). However, subcellular visualisation of B*. minutum* cells using light microscopy approaches has remained challenging mainly due to the broad spectrum autofluorescence (400–700 nm) and their relatively small cell size (Deore et al., 2024a, 2022; Nitschke et al., 2022). In our previous work, we explored fluorescence lifetime imaging microscopy (FLIM) to distinguish FISH labelled intracellular bacteria from background autofluorescence arising from *B. minutum* (Deore et al., 2024a, 2022). Although the fluorescence lifetime readout can be easily coupled with traditional confocal laser scanning microscopy, it poses an infrastructural barrier for research labs lacking sizable central research funding. Therefore, our present work attempts to overcome challenges associated with visualisation of *B. minutum* using commonly used laboratory reagents (basic chemicals for polymer mesh formation, detergents, and enzymes) to remove background autofluorescence and super resolution scale imaging under diffraction limited light microscopes.

### Optical clearing of *B. minutum* autofluorescence and isotropic expansion

We hypothesized that the recently invented ExM procedure (Chen et al., 2015) would help in the optical clearing of autofluorescence chemicals in *B. minutum* because of the involvement of free-radical induced hydrogel formation and a hydration process (Richardson and Lichtman, 2015). Further, the anchoring step using GA (Chozinski et al., 2016) and AcX (Chen et al., 2015) help maintaining covalent attachment of biomolecules of interests within hydrogel. The anticipated clearance of autofluorescence is evident from the highly transparent nature of *B. minutum* cells in which optical contrast (phase contrasts images in Figures 3, 4 and 5) of the cellular features was only due to NHS ester labelling (M’Saad and Bewersdorf, 2020).Further, the line profile graphs in Figures 9a, d and b show lowered fluorescence signals in post-ExM samples compared to pre-ExM which corroborates with removal of autofluorescence.

The success of the ExM procedure is typically measured by the maximum achievable expansion factor while maintaining the overall isotropy (i.e., relative positioning of cellular features in pre- and post-ExM state). These factors largely depend on the mechanical properties of the specimen (Truckenbrodt et al., 2019), compatibility of thermo-chemical conditions (Truckenbrodt, 2023) and suitability of cell fixation treatment (Laporte et al., 2022) to preserve the ultrastructural context such that the spatial positioning between two targets is maintained relatively intact. Therefore, we strategically tweaked the classical 4-fold ExM protocol (Chen et al., 2015; Wen et al., 2021), based on knowledge of subcellular organisation of marine dinoflagellates that are taxonomically close to *B. minutum* (Kwok et al., 2023). This allowed us to achieve a maximum of 4.4-fold expansion with a circularity index of 0.9 for *B. minutum*.

### Sonication and in hydrogel enzymatic digestion are essential

Our approach firstly involved removal of the cell wall (the amphiesma) from *B. minutum* which contains thick cellulosic plates (Kwok et al., 2023) often making it recalcitrant to experimental manipulations (Pairs et al., 2024). Due to this we anticipated considerable mechanical hindrance in expansion of *B. minutum* cells. The presence of a suture-like feature in control cells (i.e., without sonication, Figure 2a) suggests that the presence of thecal plates (Kwok et al., 2023; Lau et al., 2007) possibly with intact outer plasma membrane precludes the entry of cell wall stain and other chemicals (Pairs et al., 2024). Earlier studies report of the use of chemicals such as potassium hydroxide (Levin et al., 2017) and a detergent (octyl β-D-glucopyranoside (Pairs et al., 2024), and sonication (Lau et al., 2007) for the permeabilization as well as removal of outer plasma membrane. We implemented a mild sonication step prior to the ExM procedure and observed a strong calcofluor white signal compared to control samples (Figure 2) which suggests sonication effectively removes the outer plasma membrane.

We report that the removal of the plasma membrane (through sonication) surrounding the cellulosic thecal plates possibly enhanced the effectiveness of cell wall digesting enzymes, Cellulase RS and Macerozyme R10, by improving the accessibility to cellulosic thecal plates. Although the enzymatic removal of cell wall was accompanied by reduction in autofluorescence which is in agreement with Pairs et al., 2024. In cultured plant cell material a short duration mild pre-treatment for cell wall digestion is recommended (Hawkins et al., 2023). Our observations suggests that a mild pre-treatment for shorter duration impacted the integrity (Levin et al., 2017) and dampened autofluorescence in *B. minutum* cells (Pairs et al., 2024). This is likely due to the crude nature of the enzymatic mix (extracts from mutant *Trichoderma viride*). We instead performed enzymatic cell wall digestion on *B. minutum* cells after the anchoring and gelation step (i.e., cells embedded in hydrogel) without compromising overall integrity of cell shape which is likely due to strong covalent bonding of cellular material with the hydrogel matrix.

A few variant methods of ExM utilise a single step approach for fixation and biomacromolecule anchoring using combinations of FA and AA for subcellular visualisation of protists (Mikus et al., 2025) including malaria parasite (Bertiaux et al., 2021), microalgae (Laporte et al., 2022) and yeast (Chen et al., 2021). However, we observed that a separate PFA-mediated cell fixation followed by biomolecular anchoring using GA yields resulted in homogenous and consistent non-specific (cytoplasmic protein) as well as specific (F-actin, calmodulin, 18S rRNA) post-ExM labelling in *B. minutum* (Figures 3 and 9). Moreover, the preservation of an intact spherical pyrenoid, as observed in Laporte et al. (2022), suggests that this ultrastructural feature of *B. minutum* is well preserved through two step approach rather than single step treatment using FA/AA. We recommend exploration of other fixation methods including cryofixation (Laporte et al., 2022) for the visualisation of other subcellular features (e.g., microtubules, endoplasmic reticulum, mitochondria, etc.,) that are not reported in this study.

### Combined thermo-chemical denaturation of proteinase K-mediated homogenisation unlocks isotropic expansion of *B. minutum*

Due to the failure in achieving expansion using either of the homogenisation treatment, proteinase K or detergent-based, we combined these two approaches. Several variant methods of ExM recommend the use SDS at 58°C for 12 h or at 95°C for 30 min (Truckenbrodt, 2023). In the case of *B. minutum*, both conditions yielded poor results, however high temperature (95°C) for relatively longer duration (3 h) showed consistent homogenisation and expansion. The combination of proteinase K after SDS-based denaturation, despite PBS washing steps in between, came with a caveat of potentially reducing the activity of the enzyme. Therefore, we introduced a specific washing regime using 1% Triton X-100 at 60°C for 20 mins (Lee et al., 1978). This washing step helps in removal of SDS which is likely to denature proteinase K added in the subsequent step. Overall, the combination of SDS-based denaturation (95°C for 3 h) followed by Triton X-100 wash (60°C for 20 mins) and proteinase K-mediated homogenisation (37°C for 3 h) consistently yielded 4-fold expansion of *B. minutum*.

### ExM compatible click chemistry tools for improved post labelling

Our approach firstly involved expansion of *B. minutum* cells pre-labelled using FISH oligonucleotide targeting 18S rRNA through linker 2 which yielded very weak fluorescence signals post-expansion. This linker exploits a reactive ester group for its coupling with the oligonucleotide probe and contains an acryloyl group for covalent grafting of the hybridised product in hydrogel while retaining DBCO-modified TAMRA fluorophore (Wen et al., 2021). Such reduction in fluorescence signal due to free radical chemistry of hydrogel formation is widely reported in the ExM literature (Wen et al., 2023). Therefore, we explored two different click-chemistry approaches for post-ExM labelling.

The first strategy exploits the strong affinity of biotin (FISH oligonucleotide contained biotin modification) towards streptavidin (conjugated to Alexa Fluor 488). Although we obtained strong post-ExM signals, these are likely to be confounded by the presence of intracellular bacteria (Figure S3c). *B. minutum* requires exogenous supply of biotin which is hypothesised to be provisioned by their endosymbiont bacteria (Matthews et al., 2020). Therefore, on occasions the use of biotin-streptavidin chemistry may result in false positives especially in non-axenic cultures. A second approach involves the use of separate trifunctional linker 1 containing azide reactive group interacting with DBCO-modified fluorophore Az594 added post-ExM (chemistry developed by Chrometra scientific, Belgium). Our results (Figure 9b) clearly show abundant 18S rRNA signals in the *B. minutum* cytoplasm using post-ExM labelling approaches. We therefore recommend the use of ExM compatible modified chemistries, such as oligonucleotide modification (Truckenbrodt, 2023) and trifunctional linkers (Wen et al., 2021), to covalently bind target molecule within hydrogel while enabling its detection post-ExM and maintain sufficient fluorescence signals.

The consideration of post-ExM labelling is especially important for the successful integration of FISH with ExM. This combination often comes with a trade-off in achieving the maximum expansion factor while maintaining intact oligonucleotide probe hybridisation with the target in high ionic strength buffers (Gao et al., 2017; Wen et al., 2021). The stringent conditions required for maintaining a high efficiency of the FISH procedure are inherently incompatible with the final expansion of the hydrogel in water. We overcome this critical barrier in the integration of FISH with ExM by incorporating a novel platinum (II)-based trifunctional linker 1 as described in Wen et al. (2021). This linker moiety modifies the oligonucleotide probe such that the hybridized product is covalently grafted in the hydrogel (Wen et al., 2021) and hence making it compatible with final expansion in water without the use of high ionic strength buffers. The covalently grafted FISH hybridised product is efficiently post labelled using any dibenzocyclooctyne-modified (DBCO) fluorophore without compromising fluorescence signal or expansion factor.

We explored compatibility of antibody labelling using our optimised ExM protocol however, only achieved 3-fold expansion. This is likely due to possible intereference from the seconday antibody incubation buffer containing Image-IT^TM^ FX signal enhancer (a commercial product) which is of an unknown composition. We suggest to instead use trifunctional anchors as outlined in label retention (LR-ExM) variant of the expansion procedure. These anchors contain reactive ester groups for conjugation to primary antibody, a monomer moiety participating in gelation and orthogonal reporter handles for post-ExM fluorophore labelling (Shi et al., 2021). This approach therefore will eliminate the use of secondary antibodies containing fluorophore.

### Subcellular features of *B. minutum* revealed by ExM

This work reveals subcellular features such as a pyrenoid, F-actin patches and a calcium binding protein, calmodulin, that have not previously been well distinguished in *B. minutum* cells partly due to autofluorescence (Deore et al., 2024a, 2022). Our report suggests that the pyrenoid structure is roughly spherical and surrounded by a thick protein-rich ring in *B. minutum*. We hypothesised that this proteinaceous ring surrounding pyrenoid resembles the PyShell structure described in the marine diatoms, *Phaeodactylum tricornutum* and *Thalassiosira pseudonana* (Shimakawa et al., 2024). The PyShell structure is described to be vital for the maintenance of overall architecture of the pyrenoid as well as for efficient operation of carbon concentrating mechanisms (Shimakawa et al., 2024). The exact molecular composition, function of this proteinaceous ring and importantly its presence in *B. minutum* cells *in hospite* (inside host) remains to be determined. Our ExM approach now enables spatial protein mapping of this proteinaceous ring structure potentially revealing its role.

F-actin fibres are the most widely described cytoskeletal features in the protist kingdom (Mikus et al., 2025). However, contrary to our expectations the phalloidin staining (targeting F-actin) revealed patches of actin in cytoplasmic spaces of *B. minutum*. Similar cortical actin patches are visualised in yeast, *Saccharomyces cerevisae* (Adams and Pringle, 1984) and the parasitic protozoan, *Entamoeba histolytica* (Manich et al., 2018). These cortical patches are implicated in maintaining cell polarity (Waddle et al., 1996) and interact with actin fibres thereby exerting control over cell growth (Amberg, 1998) and partake in endocytosis process (Robertson et al., 2009). *Symbiodinium kawagutti*, a closely related species to the photosymbiont used in this study, is reported to show a lattice-like actin structure with thin radiating microfibers (Villanueva et al., 2014). We solely observed actin patches with no fibrous cytoskeleton in *B. minutum*. This suggests that one of the ExM steps in this protocol is likely to be destructive to cytoskeleton fibres. Hence for the future studies, we suggest exploration of cryofixation of samples for better preservation of cytoskeletal fibres (Laporte et al. 2022), if desired. The presence of weak F-actin signals regions of proteinase ring suggests involvement of cytoskeletal features in pyrenoid positioning which co-localises with chloroplast (Craig et al., 2019). Although, non-specific cross labelling of pyrenoid ring with actin label cannot completely overruled (Craig et al., 2019). The observed abundance of 18S rRNA in *B. minutum* is in line with our expectations because of its association with 40S small ribosome subunit present in the cytoplasm of most eukaryotes (Henras et al., 2015).

The presence of substantial fluorescence surrounding *B. minutum* cells (pre-ExM panel in Figure 9e) suggests the presence of calmodulin, a calcium binding protein, across the plasma membrane which is removed by sonication prior to commencement of ExM procedure. This supports the absence of plasma membrane specific calmodulin signals in post-ExM samples (Figure 9e). Calmodulin is localised in plasma membrane of lily pollens, protoplast of tobacco (Wang et al., 2009) and *Trypanosoma equiperdum* (Ramírez-Iglesias et al., 2018), where it acts as a calcium sensor mediating cascade of cell signalising. We also report intracellular distribution of calmodulin in *B. minutum* cells (post-ExM panel in Figure 9e). This observation is corroborated by a report outlining a calcium dose dependent rise in the expression of calmodulin genes in diverse species of marine dinoflagellates such as A*mphidinium carterae*, *Cochlodinium polykrikoides*, *Prorocentrum micans*, and *P. minimum* (Abassi et al., 2021).The role of calmodulin in maintaining cellular homeostasis is reported in corals (Lock et al., 2022) however, its dynamics in *B. minutum* especially under stress conditions remains to be deciphered.

While ExM enables subcellular observation in *B. minutum* cells, the destructive nature and multi-day procedure limits its application only to fixed samples (Truckenbrodt et al., 2019). Unless thoroughly validated the possibility of spatial distortion due to anisotropic expansion of the target feature remains a major limitation (Truckenbrodt, 2023). Our ExM gel recipe is adopted from Vanheusden et al. (2020) in which isotropic expansion is confirmed using pre- and post-ExM measurement of photobleached cube generated within a cell nucleus using two-photon excitation. Despite using a pre-optimised gel recipe (Vanheusden et al., 2020; Wen et al., 2021), spatial distortions are likely to be introduced during gel handling and transfers. Therefore, it is essential to integrate intrinsic fluorescence grids (Damstra et al., 2023) to thoroughly validate isotropic expansion. Often two organelles of the same cell may require different fixation approaches to preserve their native ultrastructure (Laporte et al., 2022) and therefore, this will requires re-optimisation of the existing ExM procedure which is time intensive. Overall, our highly customised ExM protocol for *B. minutum* unlocks subcellular imaging of *B. minutum* using readily available laboratory reagents and compatible click-chemistries commercially available at fractional cost. Translation of this method to closely related coral photosymbiont species (*Cladocopium* sp., and *Durusdinium* sp.) will improve our knowledge of subcellular landscape of these microbes under stressor conditions and during stages of cell division.

## Acknowledgement

D.R.B. was supported by a Symbiosis in Aquatic Systems grant from the Gordon and Betty Moore Foundation (Grant No. 9351) and a Future Fellowship from the Australian Research Council (Grant No. FT250100040). P.D. acknowledges funding from Mary Lugton Fellowship by the University of Melbourne.

## Abbreviations

FISH: Fluorescence *in situ* hybridisation
FLIM: Fluorescence lifetime imaging microscopy
ExM: Expansion microscopy
Ex/em max: Excitation/emission maxima
DBCO: Dibenzocyclooctyne
DMSO: Dimethyl sulfoxide
GA: Glutaraldehyde
PFA: Paraformaldehyde
FA: Formaldehyde
AA: Acrylamide
BIS: N,N’-Methylene-bis-acrylamide
APS: Ammonium persulfate
TEMPO: 4-Hydroxy-2,2,6,6-tetramethylpiperidine 1-oxyl
TEMED: N,N,N’,N’-tetramethylethane-1,2-diamine
SDS: Sodium dodecyl sulfate
PG: Propyl gallate
NHS ester: N-hydroxysuccinimide esters
rRNA: Ribosomal ribonucleic acid
CLSM: Confocal laser scanning microscopy
STED: Stimulated emission depletion
fRSS: Filtered red sea salt

## References

Abassi, S., Wang, H., Ki, J.-S., 2021. Characterization and Ca2+-induced expression of calmodulin (CaM) in marine dinoflagellates. Eur. J. Protistol. 77, 125765.

Adams, A., Pringle, J.R., 1984. Relationship of actin and tubulin distribution to bud growth in wild-type and morphogenetic-mutant Saccharomyces cerevisiae. J. Cell Biol. 98, 934–945.

Amario, M., Villela, L.B., Jardim-Messeder, D., Silva-Lima, A.W., Rosado, P.M., Moura, R.L., Sachetto-Martins, G., Chaloub, R.M., Salomon, P.S., 2023. Physiological response of Symbiodiniaceae to thermal stress: Reactive oxygen species, photosynthesis, and relative cell size. PLoS One 18, 0284717.

Amberg, D.C., 1998. Three-dimensional imaging of the yeast actin cytoskeleton through the budding cell cycle. Mol. Biol. Cell 9, 3259–3262.

Bertiaux, E., Balestra, A.C., Bournonville, L., Louvel, V., Maco, B., Soldati-Favre, D., Brochet, M., Guichard, P., Hamel, V., 2021. Expansion microscopy provides new insights into the cytoskeleton of malaria parasites including the conservation of a conoid. PLoS Biol. 19, 3001020.

Camp, E.F., Kahlke, T., Nitschke, M.R., Varkey, D., Fisher, N.L., Fujise, L., Goyen, S., Hughes, D.J., Lawson, C.A., Ros, M., 2020. Revealing changes in the microbiome of Symbiodiniaceae under thermal stress. Environ. Microbiol. 22, 1294–1309.

Chen, F., Tillberg, P.W., Boyden, E.S., 2015. Expansion microscopy. Science 347, 543–548.

Chen, L., Yao, L., Zhang, L., Fei, Y., Mi, L., Ma, J., 2021. Applications of super resolution expansion microscopy in yeast. Front. Phys. 9, 650353.

Chozinski, T.J., Halpern, A.R., Okawa, H., Kim, H.-J., Tremel, G.J., Wong, R.O., Vaughan, J.C., 2016. Expansion microscopy with conventional antibodies and fluorescent proteins. Nat. Methods 13, 485–488.

Craig, E.W., Mueller, D.M., Bigge, B.M., Schaffer, M., Engel, B.D., Avasthi, P., 2019. The elusive actin cytoskeleton of a green alga expressing both conventional and divergent actins. Mol. Biol. Cell 30, 2827–2837.

Damstra, H.G., Passmore, J.B., Serweta, A.K., Koutlas, I., Burute, M., Meye, F.J., Akhmanova, A., Kapitein, L.C., 2023. GelMap: intrinsic calibration and deformation mapping for expansion microscopy. Nat. Methods 20, 1573–1580.

Deore, P., Ching, S.J.T.M., Brumley, D.R., van Oppen, M.J., Hinde, E., Blackall, L.L., 2024a. Cutting through host autofluorescence: fluorescence lifetime imaging microscopy for visualising intracellular bacteria in Symbiodiniaceae. bioRxiv.

Deore, P., Ching, S.J.T.M., Nitschke, M.R., Rudd, D., Brumley, D.R., Hinde, E., Blackall, L.L., van Oppen, M.J., 2024b. Unique photosynthetic strategies employed by closely related Breviolum minutum strains under rapid short-term cumulative heat stress. J. Exp. Bot. 75, 4005–4023.

Deore, P., Wanigasuriya, I., Ching, S.J.T.M., Brumley, D.R., van Oppen, M.J., Blackall, L.L., Hinde, E., 2022. Fluorescence lifetime imaging microscopy (FLIM): a non-traditional approach to study host-microbial symbioses. Microbiol. Aust. 43, 22–27.

Gambarotto, D., Hamel, V., Guichard, P., 2021. Ultrastructure expansion microscopy (U-ExM. Methods Cell Biol. Elsevier 161, 57–81.

Gao R., Asano S.M., Boyden E.S., 2017. Q&A: expansion microscopy. BMC Biol. 15, 1–9.

Gornik, S.G., Maegele, I., Hambleton, E.A., Voss, P.A., Waller, R.F., Guse, A., 2022. Nuclear transformation of a dinoflagellate symbiont of corals. Front. Mar. Sci. 9, 1035413.

Hawkins, T.J., Robson, J.L., Cole, B., Bush, S.J., 2023. Expansion Microscopy of Plant Cells (PlantExM). The Plant Cytoskeleton: Methods and Protocols. Springer US, New York, NY.

Henras, A.K., Plisson-Chastang, C., O’Donohue, M.F., Chakraborty, A., Gleizes, P.E., 2015. An overview of pre-ribosomal RNA processing in eukaryotes. Wiley Interdiscip. Rev. RNA 6, 225–242.

Hughes, T.P., Anderson, K.D., Connolly, S.R., Heron, S.F., Kerry, J.T., Lough, J.M., Baird, A.H., Baum, J.K., Berumen, M.L., Bridge, T.C., 2018. Spatial and temporal patterns of mass bleaching of corals in the Anthropocene. Science 359, 80–83.

Ishii, Y., Ishii, H., Kuroha, T., Yokoyama, R., Deguchi, R., Nishitani, K., Minagawa, J., Kawata, M., Takahashi, S., Maruyama, S., 2023. Environmental pH signals the release of monosaccharides from cell wall in coral symbiotic alga. eLife 12, 80628.

Jinkerson, R.E., Russo, J.A., Newkirk, C.R., Kirk, A.L., Chi, R.J., Martindale, M.Q., Grossman, A.R., Hatta, M., Xiang, T., 2022. Cnidarian-Symbiodiniaceae symbiosis establishment is independent of photosynthesis. Curr. Biol. 32, 2402–2415. e4.

Kirk, A.L., Clowez, S., Lin, F., Grossman, A.R., Xiang, T., 2020. Transcriptome reprogramming of Symbiodiniaceae Breviolum minutum in response to casein amino acids supplementation. Front. Physiol. 11, 574654.

Kwok, A.C.M., Chan, W.S., Wong, J.T.Y., 2023. Dinoflagellate amphiesmal dynamics: Cell wall deposition with ecdysis and cellular growth. Mar. Drugs 21, 70.

Laporte, M.H., Klena, N., Hamel, V., Guichard, P., 2022. Visualizing the native cellular organization by coupling cryofixation with expansion microscopy (Cryo-ExM). Nat. Methods 19, 216–222.

Lau, R.K., Kwok, A., Chan, W., Zhang, T., Wong, J.T., 2007. Mechanical characterization of cellulosic thecal plates in dinoflagellates by nanoindentation. J. Nanosci. Nanotechnol. 7, 452–457.

Lee, L., Deas, J., Howe, C., 1978. Removal of unbound sodium dodecyl sulfate (SDS) from proteins in solution by electrophoresis through Triton X-100-agarose. J. Immunol. Methods 19, 69–75.

Levin, R.A., Suggett, D.J., Nitschke, M.R., van Oppen, M.J., Steinberg, P.D., 2017. Expanding the Symbiodinium (Dinophyceae, Suessiales) toolkit through protoplast technology. J. Eukaryot. Microbiol. 64, 588–597.

Lima, M.S., Hamerski, L., Silva, T.A., Cruz, M.L.R., Varasteh, T., Tschoeke, D.A., Atella, G.C., Souza, W., Thompson, F.L., Thompson, C.C., 2022. Insights on the biochemical and cellular changes induced by heat stress in the Cladocopium isolated from coral Mussismilia braziliensis. Front. Microbiol. 13, 973980.

Lock, C., Bentlage, B., Raymundo, L.J., 2022. Calcium homeostasis disruption initiates rapid growth after micro-fragmentation in the scleractinian coral Porites lobata. Ecol. Evol. 12, 9345.

Maire, J., Girvan, S.K., Barkla, S.E., Perez-Gonzalez, A., Suggett, D.J., Blackall, L.L., van Oppen, M.J., 2021. Intracellular bacteria are common and taxonomically diverse in cultured and in hospite algal endosymbionts of coral reefs. ISME J. 15, 2028–2042.

Majerová, E., Drury, C., 2022. Thermal preconditioning in a reef-building coral alleviates oxidative damage through a BI-1-mediated antioxidant response. Front. Mar. Sci. 9, 971332.

Manich, M., Hernandez-Cuevas, N., Ospina-Villa, J.D., Syan, S., Marchat, L.A., Olivo-Marin, J.-C., Guillén, N., 2018. Morphodynamics of the actin-rich cytoskeleton in Entamoeba histolytica. Front. Cell. Infect. Microbiol. 8, 179.

Matthews, J.L., Raina, J.B., Kahlke, T., Seymour, J.R., van Oppen, M.J., Suggett, D.J., 2020. Symbiodiniaceae-bacteria interactions: rethinking metabolite exchange in reef-building corals as multi-partner metabolic networks. Environ. Microbiol. 22, 1675–1687.

Mikus, F., Ramos, A.R., Shah, H., Hellgoth, J., Olivetta, M., Borgers, S., Saint-Donat, C., Araújo, M., Bhickta, C., Cherek, P., 2025. Charting the landscape of cytoskeletal diversity in microbial eukaryotes. Cell 188, 7610–7628 7613.

M’Saad, O., Bewersdorf, J., 2020. Light microscopy of proteins in their ultrastructural context. Nat. Commun. 11, 3850.

Nitschke, M.R., Rosset, S.L., Oakley, C.A., Gardner, S.G., Camp, E.F., Suggett, D.J., Davy, S.K., 2022. The diversity and ecology of Symbiodiniaceae: A traits-based review. Adv. Mar. Biol. 92, 55–127.

Oakley, C.A., Pontasch, S., Fisher, P.L., Wilkinson, S.P., Keyzers, R.A., Krueger, T., Dove, S., Hoegh-Guldberg, O., Leggat, W., Davy, S.K., 2022. Thylakoid fatty acid composition and response to short-term cold and heat stress in high-latitude Symbiodiniaceae. Coral Reefs 41, 343–353.

Pairs, P.I., Dundon, M.L., Narváez-Vásquez, J., Orozco-Cárdenas, M.L., Xiang, T., Jinkerson, R.E., Rao, M.P., 2024. Cell wall digestion of the dinoflagellate Breviolum minutum. J. Appl. Phycol. 36, 181–189.

Pasaribu, B., Li, Y.-S., Kuo, P.-C., Lin, I.-P., Tew, K.S., Tzen, J.T., Liao, Y.K., Chen, C.-S., Jiang, P.-L., 2016. The effect of temperature and nitrogen deprivation on cell morphology and physiology of Symbiodinium. Oceanologia 58, 272–278.

Ramírez-Iglesias, J.R., Pérez-Gordones, M.C., Castillo, J.R.D., Mijares, A., Benaim, G., Mendoza, M., 2018. Identification and characterization of a calmodulin binding domain in the plasma membrane Ca2+-ATPase from Trypanosoma equiperdum. Mol. Biochem. Parasitol. 222, 51–60.

Richardson, D.S., Lichtman, J.W., 2015. Clarifying tissue clearing. Cell 162, 246–257.

Robertson, A.S., Smythe, E., Ayscough, K.R., 2009. Functions of actin in endocytosis. Cell. Mol. Life Sci. 66, 2049–2065.

Rosic, N., Delamare-Deboutteville, J., Dove, S., 2024. Heat stress in symbiotic dinoflagellates: Implications on oxidative stress and cellular changes. Sci. Total Environ. 944, 173916.

Schindelin, J., Arganda-Carreras, I., Frise, E., Kaynig, V., Longair, M., Pietzsch, T., Preibisch, S., Rueden, C., Saalfeld, S., Schmid, B., 2012. Fiji: an open-source platform for biological-image analysis. Nat. Methods 9, 676–682.

Shi, X., Li, Q., Dai, Z., Tran, A.A., Feng, S., Ramirez, A.D., Lin, Z., Wang, X., Chow, T.T., Chen, J., 2021. Label-retention expansion microscopy. J. Cell Biol. 220, 202105067.

Shimakawa, G., Demulder, M., Flori, S., Kawamoto, A., Tsuji, Y., Nawaly, H., Tanaka, A., Tohda, R., Ota, T., Matsui, H., 2024. Diatom pyrenoids are encased in a protein shell that enables efficient CO2 fixation. Cell 187, 5919–5934 5919.

Slavov, C., Schrameyer, V., Reus, M., Ralph, P.J., Hill, R., Büchel, C., Larkum, A.W., Holzwarth, A.R., 2016. “Super-quenching” state protects Symbiodinium from thermal stress—implications for coral bleaching. Biochim. Biophys. Acta BBA-Bioenerg. 1857, 840–847.

Sun, D. -e, Fan, X., Shi, Y., Zhang, H., Huang, Z., Cheng, B., Tang, Q., Li, W., Zhu, Y., Bai, J., 2021. Click-ExM enables expansion microscopy for all biomolecules. Nat. Methods 18, 107–113.

Tchernov, D., Gorbunov, M.Y., Vargas, C., Yadav, S.N., Milligan, A.J., Häggblom, M., Falkowski, P.G., 2004. Membrane lipids of symbiotic algae are diagnostic of sensitivity to thermal bleaching in corals. Proc. Natl. Acad. Sci. 101, 13531–13535.

Tortorelli, G., Rosset, S.L., Sullivan, C.E., Woo, S., Johnston, E.C., Walker, N.S., Hancock, J.R., Caruso, C., Varela, A.C., Hughes, K., 2025. Heat-induced Stress Modulates Cell Surface Glycans and Membrane Lipids of Coral Symbionts. ISME J. wraf073.

Truckenbrodt, S., 2023. Expansion microscopy: super-resolution imaging with hydrogels. Anal. Chem. 95, 3–32.

Truckenbrodt, S., Sommer, C., Rizzoli, S.O., Danzl, J.G., 2019. A practical guide to optimization in X10 expansion microscopy. Nat. Protoc. 14, 832–863.

Valdes, P.A., Yu, C.-C., Aronson, J., Ghosh, D., Zhao, Y., An, B., Bernstock, J.D., Bhere, D., Felicella, M.M., Viapiano, M.S., 2024. Improved immunostaining of nanostructures and cells in human brain specimens through expansion-mediated protein decrowding. Sci. Transl. Med. 16, 0049.

van Oppen, M.J., Raina, J.B., 2022. Coral holobiont research needs spatial analyses at the microbial scale. Environ. Microbiol. 25, 179.

Vanheusden, M., Vitale, R., Camacho, R., Janssen, K.P., Acke, A., Rocha, S., Hofkens, J., 2020. Fluorescence photobleaching as an intrinsic tool to quantify the 3D expansion factor of biological samples in expansion microscopy. ACS Omega 5, 6792–6799.

Villanueva, M.A., Arzápalo-Castañeda, G., Castillo-Medina, R.E., 2014. The actin cytoskeleton organization and disorganization properties of the photosynthetic dinoflagellate Symbiodinium kawagutii in culture. Can. J. Microbiol. 60, 767–775.

Waddle, J.A., Karpova, T.S., Waterston, R.H., Cooper, J.A., 1996. Movement of cortical actin patches in yeast. J. Cell Biol. 132, 861–870.

Wang, Q., Chen, B., Liu, P., Zheng, M., Wang, Y., Cui, S., Sun, D., Fang, X., Liu, C.-M., Lucas, W.J., 2009. Calmodulin binds to extracellular sites on the plasma membrane of plant cells and elicits a rise in intracellular calcium concentration. J. Biol. Chem. 284, 12000–12007.

Wassie, A.T., Zhao, Y., Boyden, E.S., 2019. Expansion microscopy: principles and uses in biological research. Nat. Methods 16, 33–41.

Wen, G., Leen, V., Rohand, T., Sauer, M., Hofkens, J., 2023. Current progress in expansion microscopy: chemical strategies and applications. Chem. Rev. 123, 3299–3323.

Wen, G., Vanheusden, M., Leen, V., Rohand, T., Vandereyken, K., Voet, T., Hofkens, J., 2021. A universal labeling strategy for nucleic acids in expansion microscopy. J. Am. Chem. Soc. 143, 13782–13789.

Yokouchi, H., Takeyama, H., Miyashita, H., Maruyama, T., Matsunaga, T., 2003. In situ identification of symbiotic dinoflagellates, the genus Symbiodinium with fluorescence-labeled rRNA-targeted oligonucleotide probes. J. Microbiol. Methods 53, 327–334.

