## Supplementary data for "Unlocking subcellular imaging of a cnidarian photosymbiont *Breviolum minutum*, through expansion microscopy"

**Supplementary material**

**Methods**

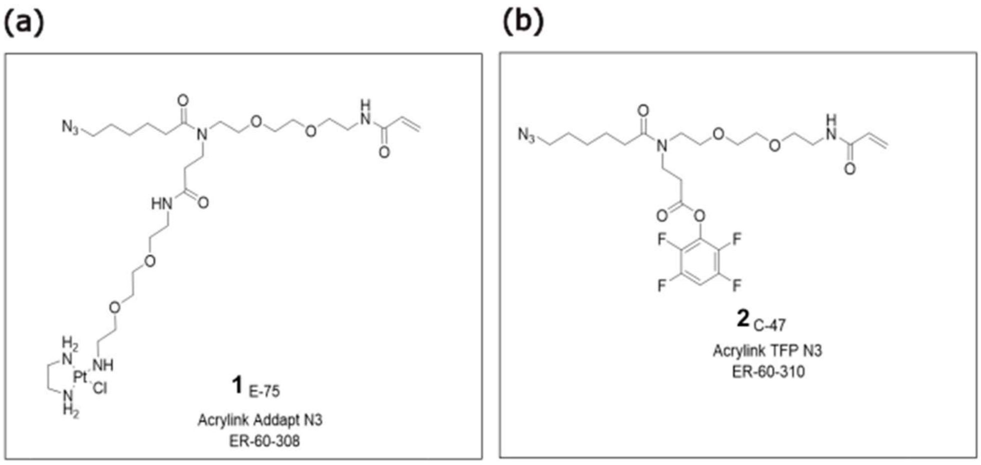


**Figure S1.** Chemical structures of tri-functional linker a) 1 and b) 2 used for covalent integration of 18S rRNA hybridised product in ExM hydrogel. The acryloyl group on the top right mimic acrylamide monomer and therefore partakes in hydrogel formation. The azide group ($–N_3$) on the top left reacts with DBCO modified fluorophores. Platinum (II) complex in linker 1 covalently bonds with N7-position of guanin nucleobase in unmodified oligonucleotide whereas, tetrafluorophenyl ester group on bottom in linker 2 reacts with primary amine group present on 5′ of the modified oligonucleotide. The descriptors underneath the label of each linker are unique catalogue identifiers provided by Chrometra Scientific, Belgium

### Results

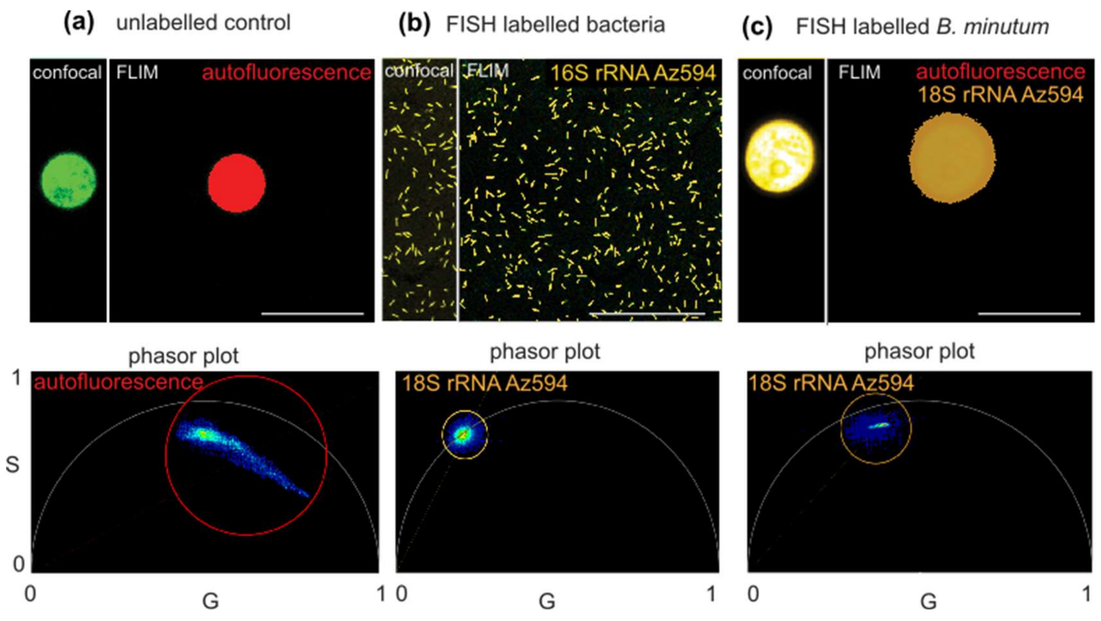


**Figure S2.** Fluorescence lifetime imaging microscopy-based (FLIM) validation for tri-functional linkers, 1 and 2, chemistries to combine fluorescence *in situ* hybridisation (FISH) with ExM for post-labelling (18S and 16S rRNA for *B. minutum* and bacteria, respectively) using DBCO modified Az594 dye.

(a) Background autofluorescence in unlabelled control *B. minutum* cell (in confocal) which was reciprocally pseudo-coloured in red (in FLIM image) using unique signature highlighted by the red circle in phasor plot. (b) Pure cultures of *Marinobacter* sp., FISH labelled using 16S rRNA oligonucleotide (EUBmix 338: 5′ GCTGCCTCCCGTAGGAGT 3′) coupled with linkers 1 or 2 (in confocal) were reciprocally pseudo-coloured in yellow (in FLIM) using unique signature highlighted by yellow coloured circle in phasor plot. Exemplar images shown in a and b serve as controls to help distinguish autofluorescence in *B. minutum* (red pseudo-coloured) from pure Az594 signals (yellow pseudo-coloured FISH labelled bacteria). (c) *B. minutum* cell FISH labelled using 18S rRNA oligonucleotide coupled with linkers1 or 2 (in confocal) was reciprocally pseudo-coloured in yellow (in FLIM) using unique signature in the phasor plot that closely resembled to pure Az594 dye signal. Autofluorescence signals were diminished as signals associated with red coloured phasor was absent in FLIM image c. Excitation laser was set to 564 nm with emission range of 600–650 nm. Threshold of 10 counts were applied for FLIM image processing which removed background interference. The phasor plot is generated through transformation of fluorescence lifetime recorded in each image pixel into a vector and represented in two-dimensional coordinate system (G and S) showing 1) the fractional contribution of exogenous fluorophores (Az594 in this case and autofluorescence, and 2) reciprocal spatial mapping of these fluorescence signals throughout source FLIM images. Scale bar=10 μm.

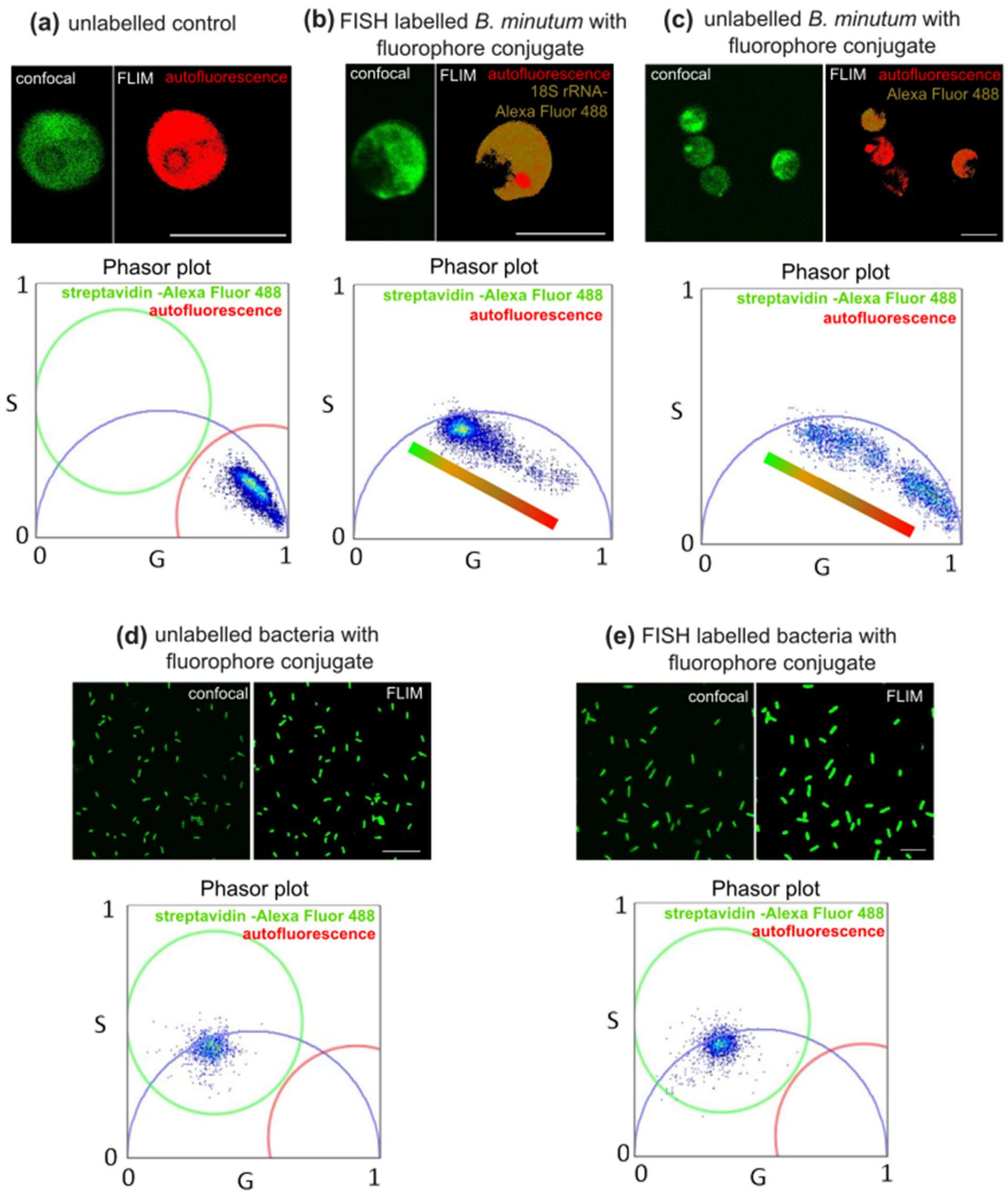


**Figure S3:** Fluorescence lifetime imaging microscopy-based (FLIM) validation for chemically modified fluorescence *in situ* hybridisation (FISH) oligonucleotides with acrydite on 5′ and biotin 3′ (18S and 16S rRNA for *B. minutum* and bacteria, respectively) post labelled using streptavidin conjugate containing Alexa Fluor 488.

(a) Background autofluorescence in an unlabelled control *B. minutum* cell (in confocal) which was reciprocally pseudo-coloured in red (in FLIM image) using the unique signature highlighted by the red coloured circle in the phasor plot. (b) A *B. minutum* cell FISH labelled using chemically modified 18S rRNA oligonucleotide (in confocal) was reciprocally pseudo-coloured in mixed green-to-red gradient (in FLIM) using the signature in the phasor plot. (c) Unlabelled *B. minutum* cells (in confocal) treated with streptavidin conjugated to Alexa Fluor 488 (negative control for 18S rRNA) were reciprocally pseudo-coloured in mixed green-to-red gradient (in FLIM) using the signature in the phasor plot. Green coloured signals indicate fluorescence from streptavidin conjugate with Alexa Fluor 488 and red coloured signals represent autofluorescence. The pixels containing mixed signals from Alexa Fluor 488 and autofluorescence form a gradient of green and red. (d) Unlabelled pure cultures of *Marinobacter* sp., treated with streptavidin conjugated to Alexa Fluor 488 (negative control for 16S rRNA) were reciprocally pseudo-coloured in mixed green-to-red gradient (in FLIM) using the signature in the phasor plot. (e) Pure cultures of *Marinobacter* sp., FISH labelled (in confocal) using chemically modified 16S rRNA oligonucleotide (EUBmix 338: 5′ GCTGCCTCCCGTAGGAGT 3′) probe and were reciprocally pseudo-coloured (in FLIM) using unique signature highlighted by green coloured circle in phasor plot. Exemplar images shown in a and d serve as controls to help distinguish autofluorescence in *B. minutum* (red pseudo-coloured) from Alexa Fluor 488 (green pseudo-coloured FISH labelled bacteria) signals. The excitation laser was set to 488 nm with an emission range of 510–550 nm. A threshold of 10 counts was applied for FLIM image processing which removed background interference. The phasor plot was generated through transformation of fluorescence lifetime recorded in each image pixel into a vector and represented in two-dimensional coordinate system (G and S) showing 1) the fractional contribution of exogenous fluorophores (Alexa Fluor 488 in this case and autofluorescence, and 2) the reciprocal spatial mapping of these fluorescence signals throughout source FLIM images. Scale bar=10 μm.

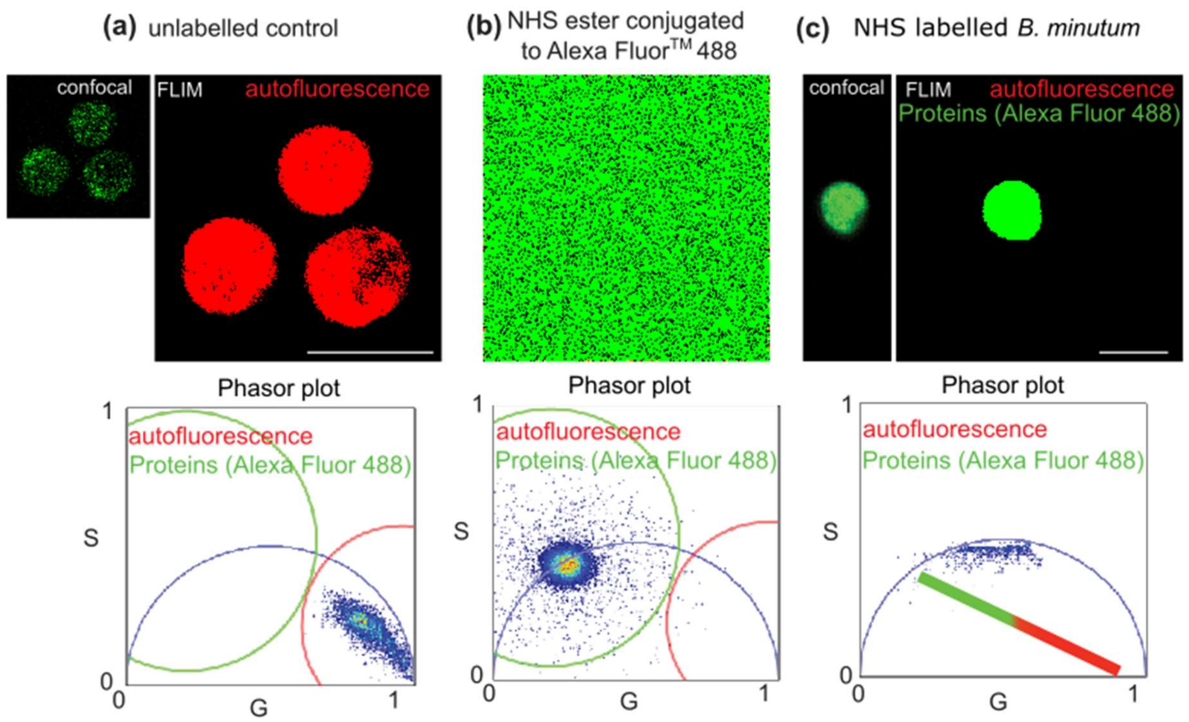


**Figure S4.** Fluorescence lifetime imaging microscopy-based (FLIM) validation for non-specific labelling of proteins using NHS ester coupled to Alexa Fluor 488 in *B. minutum*.

(a) Background autofluorescence in unlabelled control *B. minutum* cells (in confocal) which was reciprocally pseudo-coloured in red (in FLIM image) using the unique signature highlighted by the red coloured circle in the phasor plot. (b) Signals from pure NHS ester coupled to Alexa Fluor 488™ was reciprocally pseudo-coloured in green using the unique signature highlighted by the green coloured circle in the phasor plot. (c) Signals for non-specifically labelled proteins (in confocal) in *B. minutum* cells are reciprocally pseudo-coloured in green (in FLIM images) because of their close resemblance to pure Alexa Fluor 488™ signals in the phasor plot b. Autofluorescence signals were diminished as signals associated with the red coloured phasor were absent in FLIM image c). Green-to-red coloured gradient palette indicates FLIM signal arising from Alexa Fluor 488 and autofluorescence signals in *B. minutum*. The excitation laser was set to 488 nm with an emission range of 510–550 nm. A threshold of 10 counts was applied for FLIM image processing which removed background interference. The phasor plot was generated through transformation of fluorescence lifetime recorded in each image pixel into a vector and represented in two-dimensional coordinate system (G and S) showing 1) the fractional contribution of exogenous fluorophores (Az594 in this case and autofluorescence, and 2) the reciprocal spatial mapping of these fluorescence signals throughout source FLIM images. Scale bar=10 μm.

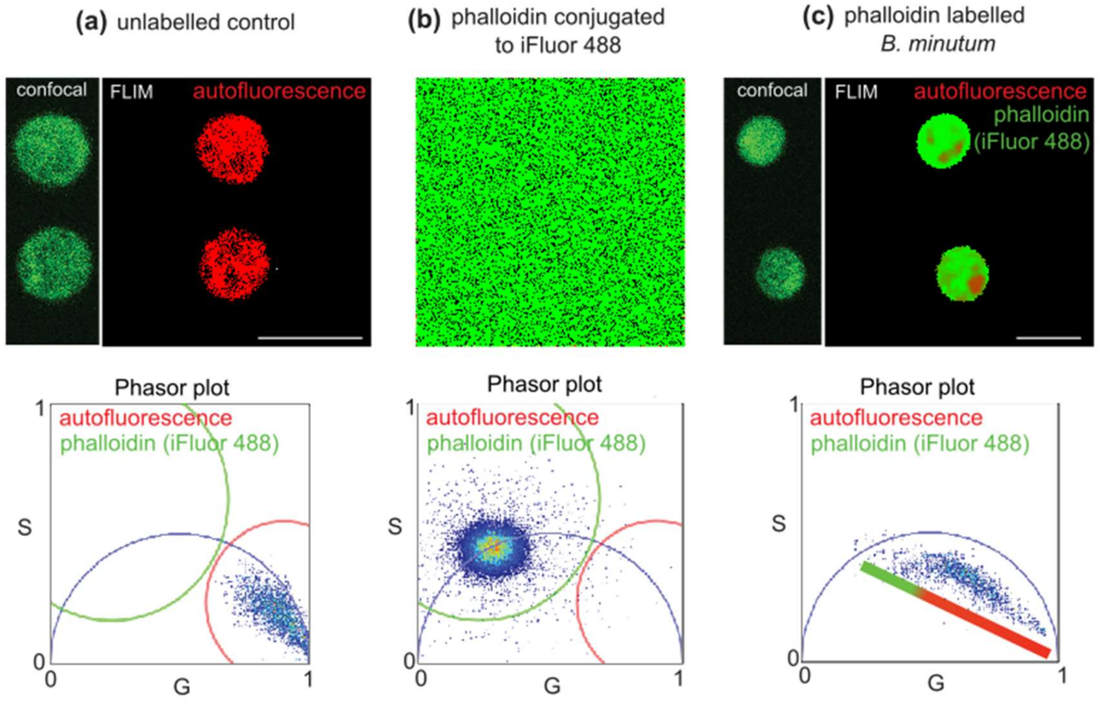


**Figure S5.** Fluorescence lifetime imaging microscopy-based (FLIM) validation for F-actin labelling using phalloidin conjugated with iFlour 488 in *B. minutum*.

(a) Background autofluorescence in unlabelled control *B. minutum* cells (in confocal) which was reciprocally pseudo-coloured in red (in FLIM image) using unique signature highlighted by red coloured circle in phasor plot. (b) Signals from pure iFluor 488 was reciprocally pseudo-coloured in green using unique signature highlighted by green coloured circle in phasor plot. (c) Signals for F-actin (in confocal) in *B. minutum* cells is reciprocally pseudo-coloured in green (in FLIM images) because of its close resemblance to pure iFluor 488 signals observed in phasor plot b. FLIM modality clearly distinguishes background autofluorescence (pseudo-coloured in red) from F-actin signals (pseudo-coloured in green) in *B. minutum* cells labelled with phalloidin in panel b. Green to red coloured gradient palette indicate FLIM signal from phalloidin (F-actin) and autofluorescence signals in *B. minutum*. Excitation laser was set to 488 nm with emission range of 500–550 nm. Threshold of 10 counts were applied for FLIM image processing which removed background interference. Phasor plot is generated through transformation of fluorescence lifetime recorded in each image pixel into a vector and represented in two-dimensional coordinate system (G and S) showing 1) the fractional contribution of exogenous fluorophores (Az594 in this case and autofluorescence, and 2) reciprocal spatial mapping of these fluorescence signals throughout source FLIM images. Scale bar=10 μm.

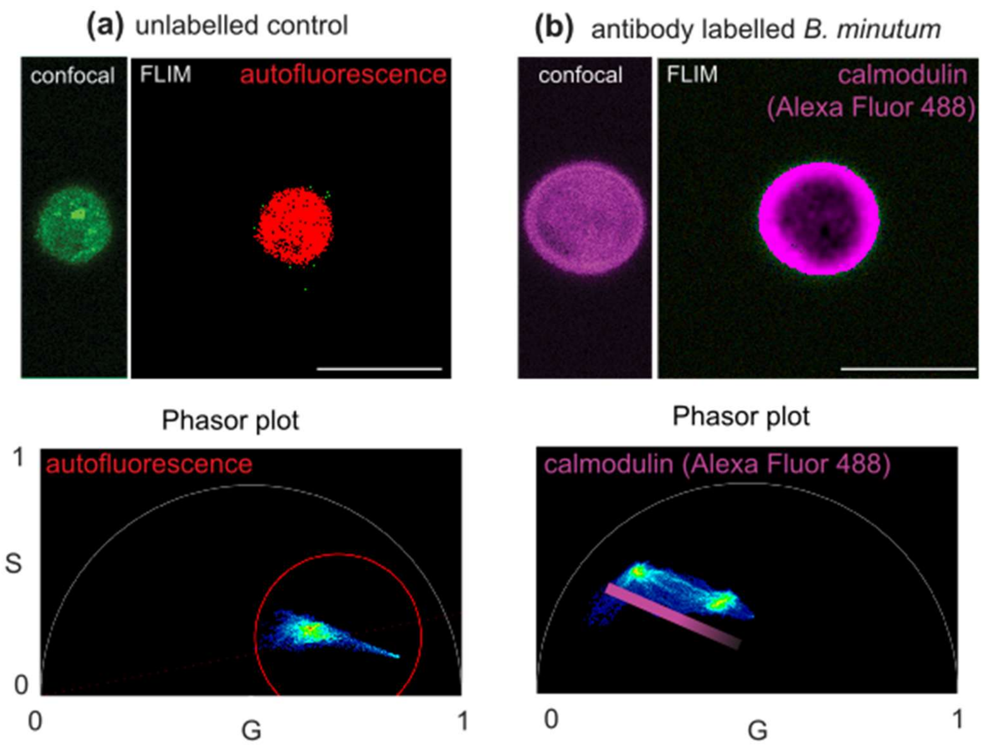


**Figure S6.** Fluorescence lifetime imaging microscopy-based (FLIM) validation for antibody labelling of calcium binding protein, calmodulin, in *B. minutum*.

(a) Background autofluorescence in unlabelled control *B. minutum* cell (in confocal) which was reciprocally pseudo-coloured in red (in FLIM image) using unique signature highlighted by red coloured circle in phasor plot. (b) *B. minutum* cell labelled using monoclonal antibody 6D4 for calmodulin protein which was detected using anti mouse secondary antibody coupled with Alexa Fluor$^{TM}$ 488 (in confocal). The unique phasor signature of Alexa Fluor 488$^{TM}$ as shown in Figure S3 was used to distinguish *B. minutum* autofluorescence from calmodulin specific (magenta coloured) signals. Intense to faded magenta coloured gradient palette indicate FLIM signal gradient of calmodulin and autofluorescence signals in *B. minutum*. Excitation laser was set to 488 nm with emission range of 510–550 nm. Threshold of 10 counts were applied for FLIM image processing which removed background interference. Phasor plot is generated through transformation of fluorescence lifetime recorded in each image pixel into a vector and represented in two-dimensional coordinate system (G and S) showing 1) the fractional contribution of exogenous fluorophores (Az594 in this case and autofluorescence, and 2) reciprocal spatial mapping of these fluorescence signals throughout source FLIM images. Scale bar=10 μm.

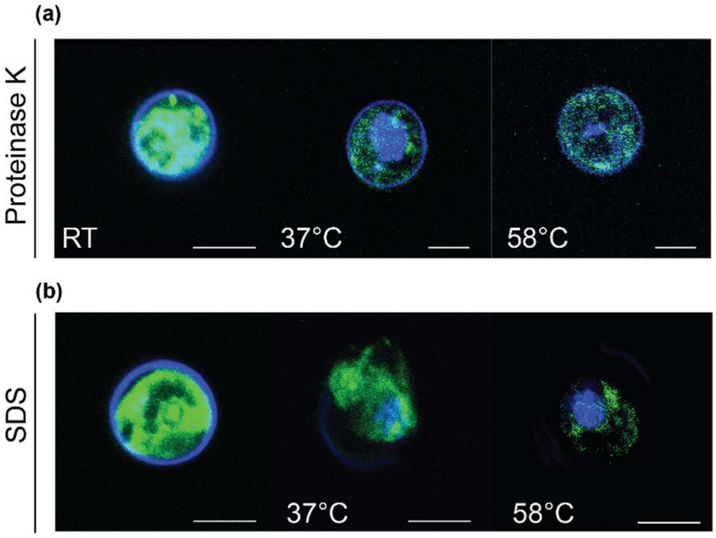


**Figure S7.** Single step approach involving either the usage of (a) classical proteinase K-mediated homogenisation or (b) sodium dodecyl sulphate (SDS) based denaturation of *B. minutum* cells at 37 and 58°C. All images are a superimposition of green (indicating autofluorescence and proteins labelled using NHS ester Alexa Fluor 488) and blue (indicating cell wall stained using calcofluor white and nucleic acid using DAPI) channels. Scale bars = 10 µm.

### Tables

**Table S1**: Increase in the mean diameter of post-ExM compared to pre-ExM and expansion factor (EF) of *B. minutum* cells under a range of thermochemical and enzymatic modifications. Welch two sample t-test summaries statistical differences with the level of significance of 0.05.

| | **pre-ExM** | | **post-ExM** | | **EP** | **t-test** | | |
|---|---|---|---|---|---|---|---|---|
| **Experimetal modifications** | **Mean diameter (µm)** | **n** | **Mean diameter (µm)** | **n** | | **t** | **df** | ***p*-value** |
| Single step proteinase K-mediated homogenisation | 9.1 | 7 | 17.3 | 3 | 2.42 | -3.23 | 4.18 | $2.99x10^{-2}$ |
| Single step SDS-based denaturation | 7.6 | 3 | 21.2 | 23 | 2.79 | -16.85 | 9.15 | $3.33x10^{-8}$ |
| Two steps shorter incubation with SDS (3h) and proteinase K (1h) | 8.2 | 11 | 24.6 | 13 | 3.0 | -35.08 | 17.70 | $2.2x10^{-16}$ |
| Two steps extended incubation with SDS and proteinase K (3h) | 7.7 | 3 | 27.4 | 23 | 3.54 | -23.32 | 22.25 | $2.2x10^{-16}$ |
| Gel recipe 2 (lower SA to AA ratio) | 7.6 | 5 | 24.5 | 8 | 3.20 | -16.45 | 7.60 | $3.2x10^{-7}$ |
| Gel recipe 1 (higher SA to AA ratio) | 8.1 | 4 | 27.9 | 18 | 3.45 | -28.47 | 19.30 | $2.2x10^{-16}$ |
| AcX anchoring | 8.2 | 4 | 26.2 | 21 | 3.16 | -22.90 | 11.28 | $8.1x10^{-11}$ |
| Separate chemical fixation (PFA) and anchoring (GA) | 7.5 | 3 | 22.8 | 6 | 3.04 | -10.9 | 5.12 | $9.6x10^{-5}$ |
| Combined cell fixation and anchoring (FA/AA) | 7.8 | 3 | 25.6 | 18 | 3.28 | -17.70 | 17.42 | $1.3x10^{-12}$ |
| ExM-antibody | 7.3 | 6 | 22.2 | 7 | 3.02 | -8.40 | 6.38 | $1.1x10^{-3}$ |
| ExM-FISH | 7.5 | 4 | 33.8 | 13 | 4.49 | -12.23 | 12.08 | $3.6x10^{-8}$ |

**Table S2**. Summary of circularity index* indicating isotropic expansion of *B. minutum* cells under a range of thermochemical and enzymatic modifications.

| **Experimetal modifications** | **Circularity index*** | |
|---|---|---|
| | **pre-ExM** | **post-ExM** |
| Single step proteinase K-mediated homogenisation | 0.90 | 0.86 |
| Single step SDS-based denaturation | 0.88 | 0.87 |
| Two steps shorter incubation with SDS (3h) and proteinase K (1h) | 0.90 | 0.86 |
| Two steps extended incubation with SDS and proteinase K (3h) | 0.92 | 0.88 |
| Gel recipe 2 (lower SA to AA ratio) | 0.96 | 0.85 |
| Gel recipe 1 (higher SA to AA ratio) | 0.90 | 0.89 |
| AcX anchoring | 0.93 | 0.91 |
| Separate chemical fixation (PFA) and anchoring (GA) | 0.96 | 0.90 |
| Combined cell fixation and anchoring (FA/AA) | 0.91 | 0.90 |
| ExM-antibody | 0.89 | 0.86 |
| ExM-FISH | 0.98 | 0.89 |

* The circularity index of 0 is considered as highly irregular shape (i.e., loss of isotropy) and 1 is considered as a perfect sphere (i.e., intact isotropy).
